# γδ T Cells Target and Ablate Senescent Cells in Aging and Alleviate Pulmonary Fibrosis

**DOI:** 10.1101/2025.05.05.652251

**Authors:** Gabriel Meca-Laguna, Tesfahun Dessale Admasu, Apoorva Shankar, Anna Barkovskaya, Isaac Collibee, Ashley Krakauer, Tommy Tran, Michael Rae, Amit Sharma

## Abstract

A variety of physiological and pathological stimuli elicit the cellular senescence response. Immune cells are known to execute surveillance of infected, cancerous, and senescent cells, and yet senescent cells accumulate with age and drive inflammation and age-related disease. Understanding the roles of different immune cells in senescent cell surveillance could enable the development of immunotherapies against biological aging and age-related disease. Here, we report the role of human gamma delta (γδ) T cells in eliminating senescent cells. Human donor Vγ9vδ2 T cells selectively remove senescent cells of different cell types and modes of induction while sparing healthy cells, with parallel findings in mouse cells. We find that senescent cells express high levels of multiple γδ T cell ligands, including cell-surface BTN3A1. Individually blocking NKG2D or γδ TCR of γδ T cells only partially reduces Vγ9vδ2 T cell cytotoxicity, evidencing their versatility in senescence removal. γδ T cells expand in response to the induction of a mouse model of idiopathic pulmonary fibrosis (IPF), accompanied by the emergence of senescent cells, and colocalize with senescent cells in lung tissue from patients with IPF. Finally, we show that adoptive cell transfer of γδ T cells into an IPF mouse model reduces the number of p21-expressing senescent cells in affected lung tissue and improves outcomes. γδ T cells or modalities that activate their surveillance activity present a potent approach for removing senescent cells and their attendant contribution to aging and disease.

## Introduction

The biological aging process is the common denominator of most human morbidity and mortality, and age-related disease and disability are becoming increasingly prevalent amid an ongoing global demographic shift (Rae et al., 2010). Significant progress has been made in understanding the underlying mechanisms of aging, including opportunities for biomedical intervention to extend life and preserve health (Campisi et al., 2019). One driver and hallmark of aging is cellular senescence, a state in which cells stably arrest the cell cycle and acquire a characteristic secretory phenotype (the senescence-associated secretory phenotype, or SASP) (Lopez-Otin et al., 2013). Senescent cells (SEN) gradually accumulate with age, contributing to tissue dysfunction and chronic local and systemic inflammation (or inflammaging) (Santoro et al., 2021). An increase in senescence burden has been extensively demonstrated to drive disease and to shorten health and lifespan (Schafer et al., 2017; Xu et al., 2018; Yousefzadeh et al., 2021). Conversely, clearance of SEN improves disease outcomes and extends lifespan in various models (Baker et al., 2016; Baker et al., 2011; Schafer et al., 2017; Wang et al., 2024). However, senescence also affects diverse physiological processes, including wound healing, preempting cancer progression, and tissue repair (de Magalhaes, 2024). How and why SEN accumulate in aging tissues is poorly understood beyond their transient involvement in such processes.

The SASP produced by SEN recruits immune cells such as macrophages, natural killer (NK), and T cells to the senescence site for their immune surveillance (Kale et al., 2020; Reen et al., 2025). Natural killer (NK) cells are believed to be the most well-investigated immune regulators of SEN (Brauning et al., 2022; Kale et al., 2020). However, their efficacy declines with aging as a result of both intrinsic and systemic changes (Brauning et al., 2022), and they are susceptible to immune evasion by SEN via mechanisms that include upregulation of the inhibitory ligand HLA-E and the cleavage of the NKG2D ligand MICA from their surface (Munoz et al., 2019; Pereira et al., 2019). Neutrophils and macrophages have also been implicated in regulating senescence burden, but their role is context-dependent, promoting pro-inflammatory or reparative responses depending on signals in the microenvironment (Kale et al., 2020). CD8^+^ T cells have been reported to recognize and selectively kill SEN. Still, as with NK cells, SEN employ mechanisms to evade CD8^+^ T cell immune clearance, including upregulation of HLA-E (Pereira et al., 2019) and immune checkpoint molecules like PD-L1/2, secretion of immunosuppressive factors (e.g., TGF-β), and modulation of the local microenvironment to suppress immune activation (Chaib et al., 2024). Additionally, invariant Natural Killer T cells (iNKT) were recently described to coordinate the removal of SEN (Arora et al., 2021). However, while iNKT cells can produce cytokines like IFN-γ and IL-4 and engage in cytotoxicity via perforin and granzyme pathways, their overall cytotoxic capacity is lower than that of NK cells or CD8^+^ T cells (Smyth et al., 2000; Wingender et al., 2010). This may limit their efficacy in clearing large or persistent populations of SEN.

There remain numerous unanswered questions about how the immune system regulates SEN, including how the regulation of SEN by immune cells varies across tissues and disease contexts, how tissue-specific signals modulate immune cell activity and SEN clearance, and how the immune system recognizes and responds to the heterogeneous phenotypes of SEN, which vary according to cell type of origin, mode of induction, and likely microenvironment and other factors (Admasu et al., 2021).

Gamma delta (γδ) T cells are a versatile subset of T lymphocytes with critical roles in innate immunity, tissue homeostasis, and disease. They are distinct from conventional alpha beta (αβ) T cells due to their T cell receptor (TCR), which consists of gamma (γ) and delta (δ) chains. The TCR repertoire of γδ T cells is less diverse than that of αβ T cells; however, unlike αβ T cells, γδ T cells can additionally be activated by a wide variety of ligands in a manner that does not require MHC antigen presentation (Deseke & Prinz, 2020). These ligands include phosphoantigens (pAgs; e.g., microbial metabolites like HMB-PP from bacteria), lipids presented by CD1 molecules, and stress-induced ligands (e.g., MICA, MICB, ULBPs). This capability enables γδ T cells to respond swiftly to infected, transformed, or stressed cells (Deseke & Prinz, 2020). γδ T cells are prevalent in epithelial-rich tissues, such as the skin (as intraepithelial lymphocytes), gut mucosa, and lungs, where they function as tissue-resident immune sentinels. They are also found in the bloodstream, although they comprise a smaller proportion of circulating T cells than αβ T cells. Because of their versatile and critical roles in immunity, tissue homeostasis, and disease, we investigated whether γδ T cells are involved in the immune surveillance of SEN.

We report that human γδ T cells (Vγ9vδ2 T cells) from multiple individuals with diverse clinical characteristics are highly selective in eliminating SEN in cell culture. We observed that γδ T cells engaged with SEN using both NKD2D and γδ TCR. Single-nucleus RNA Sequencing (snRNA-Seq.) and single-cell RNA Sequencing (scRNA-Seq.) identified high expression of γδ T cell ligands on SEN. We report a high expression of BTN3A1 on the membrane of cells induced to senescence using different insults. Mouse γδ T cells are highly cytotoxic to SEN, infiltrate the lungs after fibrotic injury, and eliminate SEN *in vivo*. Our results suggest that γδ T cells execute immune surveillance of SEN and show promise as a therapeutic intervention against age-related disease.

## Results

### γδ T cells selectively target senescent cells

As γδ T cells are highly heterogeneous, functional specialization depends on their tissue localization and TCR composition. Vγ9Vδ2 T cells are a prominent subset of γδ T cells found in humans and non-human primates. They are unique in their ability to respond to non-peptide antigens and play a critical role in bridging innate and adaptive immune responses. The TCR of Vγ9Vδ2, the subset of γδ T cells that is most abundant in the blood, is composed of a Vγ9 chain paired with a Vδ2 chain. We adopted a published method that allowed for the *ex vivo* expansion of Vγ9Vδ2 T cells using PBMCs to isolate and enrich γδ T cells from multiple human subjects (**Figure 1a**) (Kondo et al., 2008). This zoledronate (ZOL) -based expansion method has been reported to enrich γδ T cells with an effector memory phenotype and allow for a 200-fold expansion with up to 80% purity in 14 days (Kondo et al., 2008). Our FACS analysis of single, live cells confirmed significant variability in the initial proportion of cells expressing γδ TCR (γδTCR^+^CD3^+^), ranging from 1 to 10% in PBMCs from different subjects (**Supplementary Figure 1a, 1b, 1c,** left panel). Isolated PBMCs were treated with IL-2 and ZOL, which is known to increase the intracellular accumulation of isopentenyl pyrophosphate (IPP) in monocytes. Monocytes present this phosphoantigen (pAg) to γδ T cells and activate them. In cell culture, we observed an expansion of γδ T cells from 1-10% to 70-90% within 10-14 days after ZOL treatment (**Supplementary Figure 1c**, middle panel). Initially, the majority of CD3^+^ T cells in the PBMCs expressed αβ TCR (85-95%) compared to γδ TCR (5-15%) (**Figure 1b**, left panel). Following IL-2 treatment and expansion with ZOL, only about 5% of CD3^+^ cells expressed αβ TCR, while most (>90%) cells expressed γδ TCR (**Figure 1b**, middle panel). Our flow cytometric data further confirmed that, after expansion, > 80% of cells expressing γδ TCR expressed Vδ2 and not Vδ1 (**Supplementary Figure 1d, 1e**).

**Figure 1.**
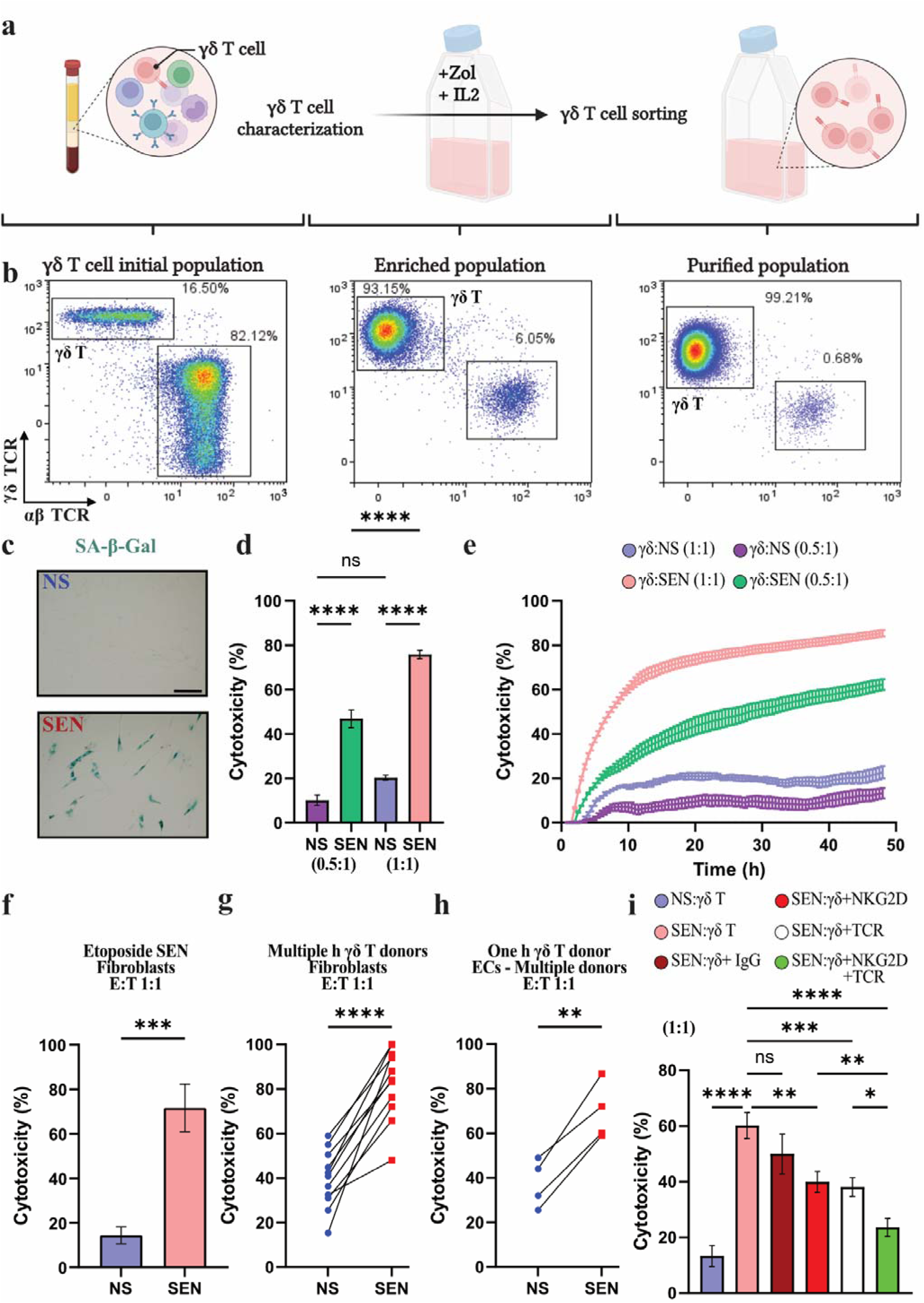
Human γδ T cells target senescent cells. (**a**) Schematic representation of the γδ T cell enrichment strategy. Peripheral blood mononuclear cells (PBMCs) were treated with Zoledronate (ZOL) and interleukin-2 (IL-2) for 24 hours and maintained in culture with IL-2 for 10-14 days leading to expansion of γδ T cells. γδ T cells were further purified using either fluorescence-activated cell sorting or magnetic-activated cell sorting. Created in https://BioRender.com. (**b**) Flow cytometry analysis of γδ T cells during enrichment. Left panel: γδ T cells identified within the total PBMC population before enrichment. Middle panel: γδ T cells following expansion and activation with ZOL and IL-2. Right panel: Highly purified γδ T cell population obtained after sorting. Representative data from experiments performed in at least three independent blood donors (n ≥ 3). (**c**) Representative images of SEN and NS IMR-90 cells after SA-β-Gal colorimetric assay. Scale bar = 150 μm. (**d**) Cytotoxicity of γδ T cells to NS controls and doxorubicin-induced SEN at E:T ratios of 0.5:1 and 1:1 after 24 hours, impedance assay. Representative data from experiments performed in at least three independent donors (n ≥ 3); one-way ANOVA. (**e**) γδ T cell 48-hour killing kinetics to NS controls and SEN at E:T ratios of 0.5:1 and 1:1. Representative data from experiments performed in at least three independent donors (n ≥ 3). (**f**) Cytotoxicity of γδ T cells to NS controls and etoposide-induced SEN after 24 hours at E:T ratio 1:1. Representative data from experiments performed in at least three independent donors (n ≥ 3); unpaired t-test. (**g**) Cytotoxicity of γδ T cells from multiple blood donors to NS controls and SEN fibroblast cells at 1:1 E:T after 24 hours. Each data point is a blood donor; unpaired t-test. (**h**) Cytotoxicity of γδ T cells from one blood donor to NS controls and SEN endothelial cells from multiple donors at 1:1 E:T. Each data point is a blood donor and represents the average of triplicates; unpaired t-test. (**i**) NKG2D and/or γδ TCR were blocked, and γδ T cells were cocultured with NS and SEN at 1:1 E:T for 24 hours. Representative data from experiments performed in at least three independent donors; one-way ANOVA. ns p > 0.05; *p ≤ 0.05; **p ≤ 0.01; ***p ≤ 0.001; ****p ≤ 0.0001.

The γδ TCR-expressing cells were purified by fluorescence- or magnetic-activated sorting to obtain >95% purity. The αβ TCR-expressing cells were a mere <5% (**Figure 1b**, right panel, **Supplementary 1c**, right panel). Further characterization of this population of γδ T cells by flow cytometry indicated that most CD3^+^ γδ TCR^+^ cells also expressed CD56 (**Supplementary Figure 1f**). CD56 (or neural cell adhesion molecule, NCAM) is a surface marker commonly associated with NK cells. It is involved in cell adhesion and immune regulation (Cooper et al., 2001), but it is also known to be expressed on γδ T cells with NK-like characteristics that are highly cytotoxic to cancer cells (Alexander et al., 2008).

We and others have shown that NK cells can selectively eliminate SEN (Kim et al., 2022). As γδ T cells exhibit innate/NK-like characteristics upon activation, enabling them to recognize stress-induced ligands on tumor cells and cells infected with pathogens displayed on SEN and elicit engagement and destruction by NK cells, we hypothesized that they may similarly surveil SEN. We used a well-characterized chemotherapy-induced senescence (CIS) cell model to test this hypothesis and generate SEN as targets. Our results confirmed a robust senescence phenotype in doxorubicin-treated human fetal lung fibroblasts (IMR-90). We observed a high number of senescence-associated β-Galactosidase (SA-β-Gal)-positive cells (**Figure 1c**), an apparent loss of nuclear HMGB1 (**Supplementary Figure 2a**), and an increase in cells with two or more persistent DNA damage foci as measured by immunofluorescence staining for γH2AX (**Supplementary Figure 2b**) compared to untreated non-senescent (NS) IMR-90 controls. Next, we conducted coculture assays with Vγ9vδ2 T cells (γδ T cells) and either SEN or NS cells using the xCELLigence RTCA instrument, which tracks cytotoxicity in real time by measuring changes in impedance that indicate the adhesion and viability of cells.

Results show minimal toxicity of γδ T cells towards NS control cells (approximately 10%), whereas 40-60% of SEN cells were killed by γδ T cells at the effector-to-target ratio (E:T) of 0.5:1, which increased to 70-80% when the E:T ratio was 1:1 after 24 hours of coculture (**Figure 1d**). Analysis of the real-time kinetics of the cytotoxicity assay indicated that γδ T targeted SEN cells within 5-10 hours after starting the coculture, with a close to maximum cytotoxicity achieved after 15-20 hours (**Figure 1e**). On the other hand, NS were minimally targeted by γδ T cells even after 48 hours of coculture (**Figure 1e**). The specificity of the cytotoxicity of γδ T cells towards SEN was further confirmed when SEN and NS IMR-90 cells were cocultured with PBMCs. We did not observe similar cytotoxicity of PBMC towards SEN or NS cells (**Supplementary Figure 2c**).

As heterogeneity of senescence is well established (Admasu et al., 2021), we investigated whether the γδ T cell-mediated cytotoxicity depends on the mode of senescence induction or the cell type of origin. Our results show that γδ T cells are highly selective in killing etoposide-treated SEN IMR-90 cells (60-80%) compared to NS cells (10-20%) at an E:T ratio of 1:1 following 24 hours in coculture (**Figure 1f**).

One potential advantage of γδ T cells as a therapeutic modality is that their cell targeting is largely MHC-independent, suggesting they are allogeneic safe. Thus, we further tested the seno-toxicity of γδ T cells from multiple donors towards SEN. γδ T cells enriched from males and females (ages 20 to 50) were significantly more toxic to SEN than NS cells and IMR-90 cells (**Figure 1g**). However, cells from some donors exhibited substantially higher toxicity towards NS cells than those from other donors (**Figure 1g**).

We reasoned that this non-specific cytotoxicity occurred because of the minor contamination of αβ T cells. Therefore, we further depleted αβ T cells from the enriched γδ T cell population. The original populations of enriched γδ T cells from donors killed 60-80% SEN cells compared to approximately 20-30% NS cells. Upon depletion of αβ T cells, γδ T cells killed SEN cells at the same rate without the depletion, but consistent with our hypothesis, toxicity towards NS cells was significantly reduced to about 0-10% (**Supplementary Figure 2d, 2e**). Our results show that γδ T cells from multiple donors can selectively kill SEN IMR-90 cells with significantly lower toxicity to NS cells following depletion of other cell types.

We next asked if γδ T cell-mediated toxicity would also be observed against SEN derived from other cell types. We induced senescence using doxorubicin treatment in human primary endothelial cells derived from the iliac artery of several donors. Following coculture with γδ T cells from an unrelated donor, we observed significantly higher toxicity towards SEN than NS cells (**Figure 1h**).

### γδ TCR and NKG2D receptors are required for the **γδ** T cell-mediated killing of senescent cells

γδ T cells use their T cell receptor (TCR) and natural killer (NK) cell-like receptors, such as NKG2D, to recognize and eliminate targets (Deseke & Prinz, 2020). To investigate how γδ T cells target SEN, we conducted a coculture experiment using the xCELLigence RTCA instrument with an effector-to-target (E:T) ratio of 1:1 between γδ T cells and SEN or NS IMR-90 cells. Our results revealed significantly higher cytotoxicity against SEN (60-80%) than NS cells. Preincubating SEN with an NKG2D neutralizing antibody before adding γδ T cells substantially decreased cytotoxicity (**Figure 1i**). A more pronounced reduction was observed when SEN cells were preincubated with a γδ TCR-neutralizing antibody (**Figure 1i**). In contrast, preincubation with isotype IgG had minimal, non-significant effects on cytotoxicity (**Figure 1i**). Preincubating with γδ TCR and NKG2D neutralizing antibodies reduced cytotoxic potential to approximately 20%. Collectively, these data suggest that γδ T cells target SEN cells through both γδ TCR and NKG2D receptors, with some functional redundancy (**Figure 1i**).

### Elevated expression of **γδ** T cell ligands on Senescent cells

To elucidate how γδ T cells recognize SEN, we performed single-nucleus RNA sequencing (snRNA-Seq.) on doxorubicin-treated SEN fibroblasts and NS control cells. Uniform manifold approximation and projection (UMAP) analysis revealed two distinct populations (**Figure 2a**). Unsupervised analysis confirmed a senescent phenotype in the doxorubicin-treated fibroblasts (**Supplementary Figure 3a**). Supervised analysis further confirmed this phenotype, showing cell cycle arrest in SEN population marked by the absence of proliferation markers such as *MKI67* or *CENPF*, and upregulation of senescence-associated pro-inflammatory genes (**Supplementary Figure 3b**). The SEN population exhibited elevated *p21* expression and lacked *TOP2A* (**Figure 2a, 2b**).

**Figure 2.**
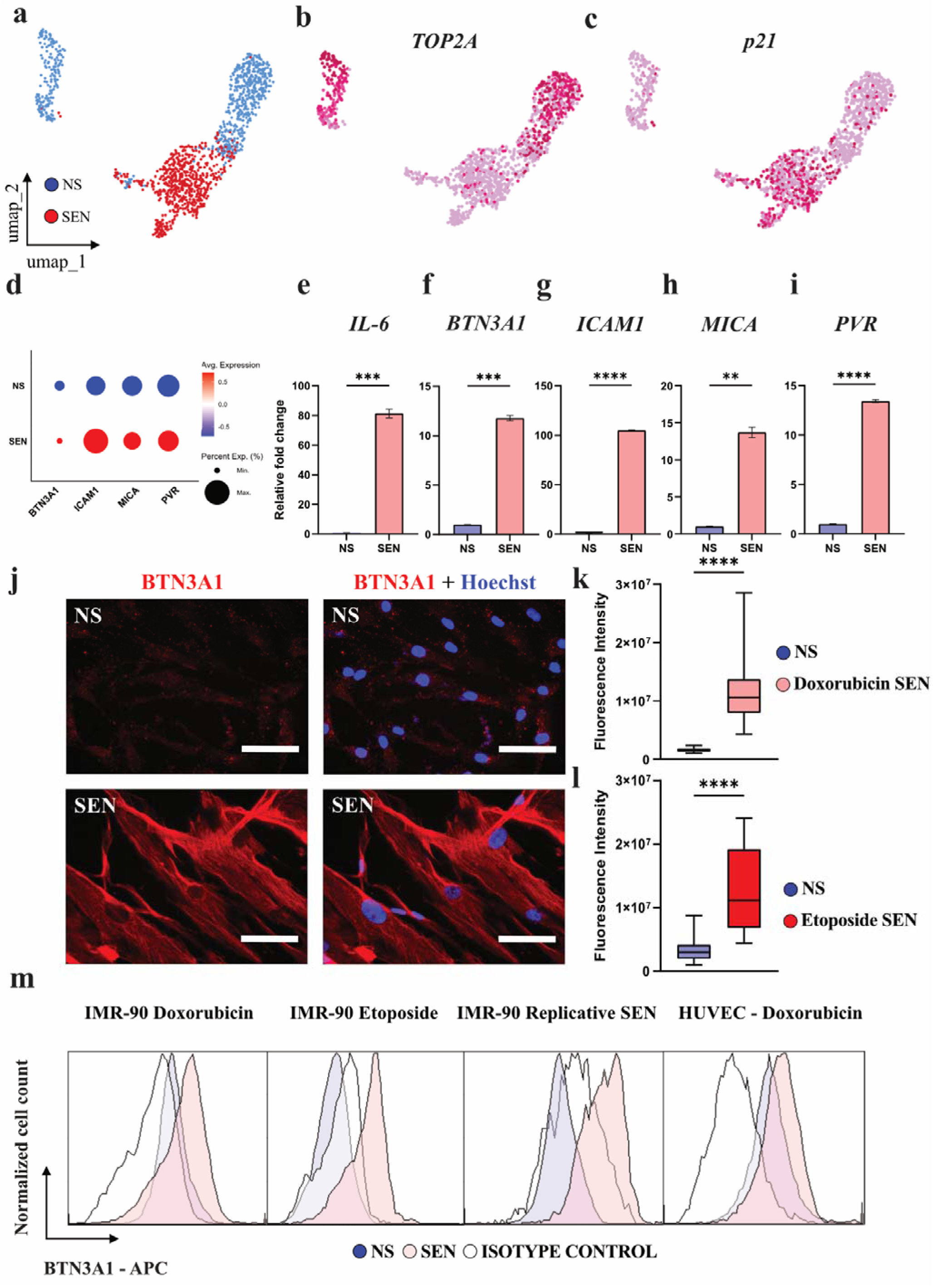
BTN3A1 and other putative γδ T cell ligands are upregulated on senescent cells. (**a**) UMAP of NS (blue) controls and doxorubicin-treated SEN (red) cells. Number of cells after quality control = 1,294 cells; NS = 668 control cells; SEN = 626 SEN. (**b**) Gene expression profile of TOP2A; dark pink represents higher gene expression. (**c**) Gene expression profile of p21; dark pink represents higher gene expression. (**d**) Gene expression dot plot of established γδ TCR ligands. (**e-i**) qPCR validation of γδ TCR ligand expression in SEN. IL-6 was used as a positive control. qPCR normalized to ACTB. Data is representative of 1 of 3 independent experiments; unpaired t-test. (**j**) Immunofluorescent membrane BTN3A1 staining of unpermeabilized NS and doxorubicin-induced SEN. BTN3A1 (red) and Hoechst (blue). Data is representative of 3 independent experiments. (**k**, **l**) Mean Fluorescence Intensity (MFI) in NS, (**k**) doxorubicin-induced SEN, and (**l**) etoposide-induced SEN. Data combined from 3 independent experiments; unpaired t-test. (**m**) Representative histograms from flow cytometry analysis of surface BTN3A1 in NS (blue), isotype control (white), and multiple SEN (light pink; doxorubicin-induced fibroblasts, doxorubicin-induced endothelial cells, replicative senescent (RS) fibroblasts, and etoposide-induced fibroblasts). Data is representative of 1 of 3 independent experiments. ns p > 0.05; *p ≤ 0.05; **p ≤ 0.01; ***p ≤ 0.001; ****p ≤ 0.0001.

γδ T cells have been previously described to recognize stress ligands on target cells, including ICAM1, PVR, BTN3A1, BTN2A1, MICA, ULBPs, NECT2, ANXA2, CD1-d, CD-1c, etc. (Deseke & Prinz, 2020). We observed elevated *ICAM1*, *PVR*, *MICA*, and *BTN3A1* expression in SEN cells. (**Figure 3d**). Quantitative RT-qPCR further confirmed a significant increase in *ICAM1*, *PVR*, *MICA*, and *BTN3A1* expression in SEN compared to NS (**Figure 3e-i**).

**Figure 3.**
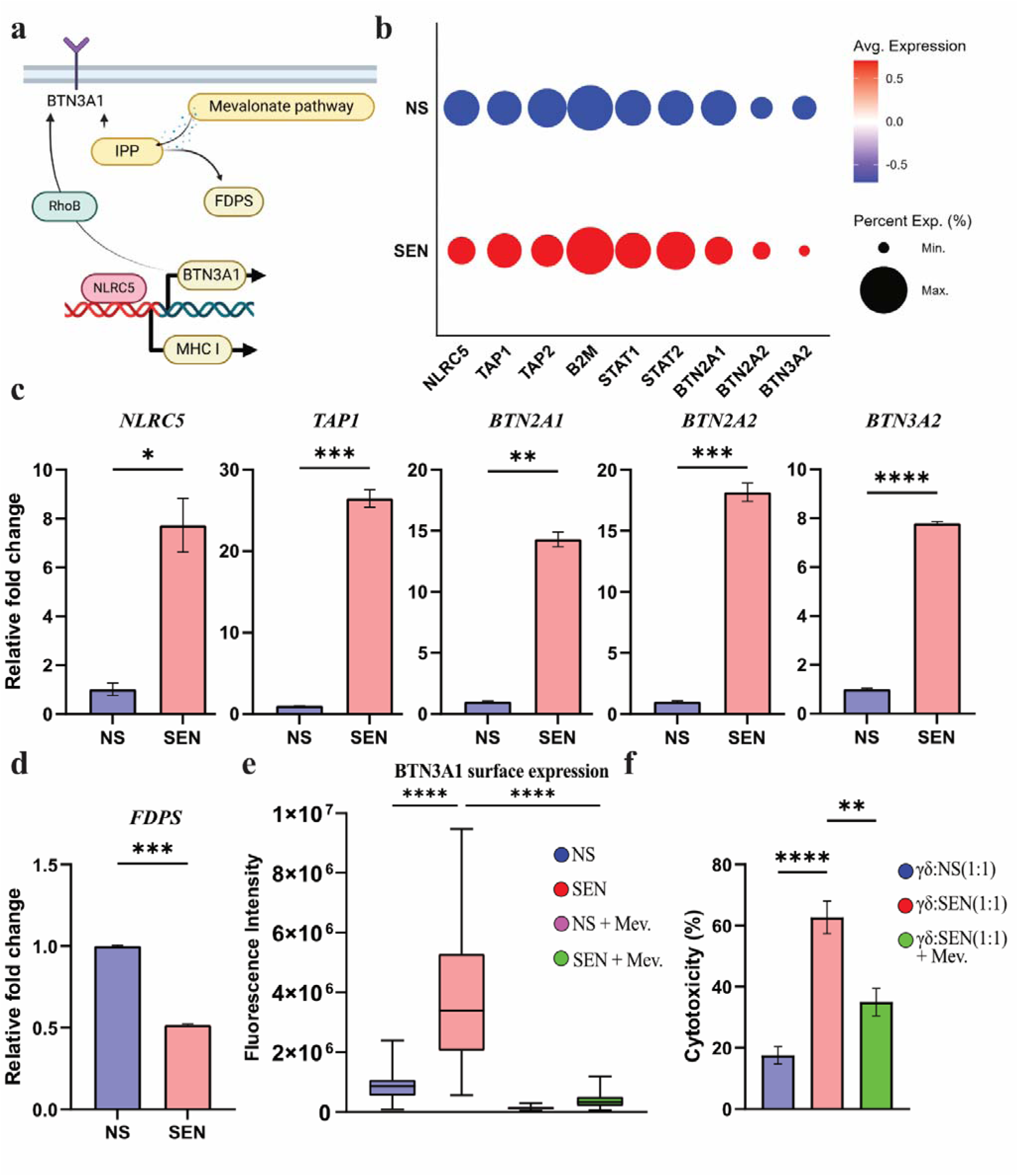
Metabolic rewiring primes senescent cell recognition by γδ T cells. (**a**) Schematic of BTN3A1 presentation on the membrane of cells targeted by γδ T cells. (**b**) Gene expression dot plot of NLRC5 and its downstream effector proteins in SEN (red) and NS (blue) controls. (**c**) qPCR validation of snRNA-Seq. Left to right, NLCR5, TAP1, BTN2A1, BTN2A2, BTN3A2 gene expression in SEN and NS. Relative fold change to ACTB. Data is representative of 1 of 3 independent experiments; unpaired t-test. (**d**) qPCR of FDPS gene expression. Relative fold change to ACTB. Data is representative of one of 3 independent experiments; unpaired t-test. (**e**) Mean fluorescence intensity of BTN3A1 surface expression with and without Mevastatin treatment. Data is representative of one of 3 independent experiments. One-way ANOVA. (**f**) Cytotoxicity of γδ T cells on SEN with or without the Mevastatin treatment. Representative data from experiments performed in at least three independent donors (n ≥ 3). One-way ANOVA. ns p > 0.05; *p ≤ 0.05; **p ≤ 0.01; ***p ≤ 0.001; ****p ≤ 0.0001.

In addition, re-analysis of two publicly available single-cell RNA Sequencing (scRNA-Seq.) datasets of SEN WI-38 cells (Wechter et al., 2023) confirmed high γδ T cell ligand expression in SEN cells (**Supplementary Figure 3c-h**). Wechter *et al*. reported the progressive establishment of senescence programming over days 1, 2, 4, 7, and 10 in WI-38 cells following etoposide treatment. Our analysis of their data showed a gradual increase in *ICAM1*, *PVR*, *MICA*, and *BTN3A1* expression after etoposide treatment in SEN (**Supplementary Figure 3c**). ScRNA-Seq. of irradiated and etoposide-treated SEN was done in this same study. A re-analysis of this data also showed increased expression of *ICAM1*, *PVR*, *MICA*, and *BTN3A1* (**Supplementary Figure 3d, 3e, 3f**). ScRNA-Seq. confirmed co-expression of γδ T cell ligands and p21 (**Supplementary Figure 3f, 3h**). These results demonstrate that SEN expresses elevated stress-associated γδ T cell ligands *ICAM1*, *PVR*, *MICA*, and *BTN3A1*.

### BTN3A1 is upregulated in the membrane of senescent cells

BTN3A1 (CD277) is a well-established ligand for γδ TCR and is increased in aggressive tumors (Payne et al., 2020). Using immunofluorescent assay in unpermeabilized cells, we confirmed higher BTN3A1 surface expression on doxorubicin- and etoposide-induced SEN fibroblasts compared to NS controls (**Figure 2j, 2k, 2l, Supplementary Figure 3i**). Mean fluorescence intensity of BTN3A1 was statistically higher in doxorubicin- and etoposide-treated SEN compared to NS controls (**Figure 2j, 2k, 2l**). Flow cytometric analysis showed a greater percentage of BTN3A1^+^ cells across multiple SEN types (doxorubicin-treated fibroblasts, doxorubicin-treated endothelial cells, replicative senescent (RS) fibroblasts, and etoposide-treated fibroblasts) (**Figure 2m**). These results suggest that the expression of BTN3A1 is elevated on the surface of SEN of multiple cell types of origin and modes of induction.

### Metabolic rewiring primes senescent cells to be recognized by **γδ** T cells

Mechanistically, γδ T cells can sense phosphoantigen accumulation through BTN3A1 (Harly et al., 2012) and BTN2A1 (Karunakaran et al., 2020). Briefly, phosphoantigens such as IPP (an intermediate metabolite of the mevalonate pathway) bind to the intracellular domain of BTN3A1, resulting in a conformational change in the extracellular domain that enables recognition by γδ TCR (Gu et al., 2017). To better understand the mechanistic regulation of BTN3A1 in SEN, we focused on its transcriptional control. NLRC5 is a key regulator of BTN3A1 expression (Mamedov et al., 2023) (**Figure 3a**). As a master regulator of MHC class I gene expression, NLRC5 acts as a transcriptional activator by binding to conserved NLRC5-responsive promoter regions (Downs et al., 2016). Our snRNA-Seq. data and quantitative real-time PCR indicate increased expression of NLRC5 and its downstream effectors BTN2A1, BTN2A2, and BTN3A2 in SEN relative to NS (**Figure 3b, 3c**).

We also observed elevated expression of several key components involved in MHC class I gene expression, such as TAP1, TAP2, B2M, STAT1, and STAT2 (**Figure 3b, 3c**). Additionally, the GTPase RhoB has been suggested to mediate and be essential for Vγ9Vδ2 T cell target recognition (Sebestyen et al., 2016). We confirmed higher expression of RhoB in SEN compared to NS controls (**Supplementary Figure 4a, 4b**). We also investigated the role of the mevalonate pathway in SEN, given its importance in biosynthesis of sterol and non-sterol isoprenoids and its potential in enhancing γδ T cell immunotherapy (Gogoi & Chiplunkar, 2013). Our quantitative real-time PCR revealed a statistically significant decrease in farnesyl diphosphate synthase (FDPS) expression in SEN (**Figure 5d**). FDPS, a downstream enzyme in the mevalonate pathway, converts IPP into farnesyl diphosphate (FPP). We hypothesized that a downregulated FDPS would lead to the accumulation of IPP in the cell, facilitating the presentation of BTN3A1 on the SEN cell surface and triggering γδ T cell engagement.

To test this, we treated SEN and NS cells with mevastatin (Mev), an inhibitor of 3-hydroxy-3-methylglutaryl-CoA reductase (HMGCR), the rate-limiting enzyme upstream of FDPS, or vehicle for 24 hours. Quantifying the immunofluorescence data indicated that treatment with Mev significantly reduced BTN3A1 expression in SEN cells but did not affect NS cells (**Figure 3e, Supplementary 4c**). Interestingly, pretreatment of SEN with 25μM Mev for 24 hours, followed by coculture with γδ T cells at an E:T ratio of 1:1, led to a 20-30% decrease in the cytotoxicity of γδ T cells toward treated SEN relative to untreated target cells (**Figure 3f**).

### Mouse **γδ** T cells target senescent cells

Given our observations where γδ T cells from multiple human blood donors selectively targeted SEN of different cell origins, we tested this same principle using mouse cells. We optimized a 7-day γδ T cell isolation, expansion, and purification strategy to expand γδ T cells 100-fold from the mouse spleen (**Figure 4a** - see Methods for details) (Williams et al., 2022). γδ T cells constitute around 1% of the total splenocyte population, which we successfully enriched and purified to >95 % (**Figure 4b, Supplementary Figure 5a, 5b, 5c**). Mouse embryonic fibroblasts (MEFs) were induced to senescence using doxorubicin. We validated a senescent phenotype in MEFs (**Figure 4c**). γδ T cells selectively targeted SEN doxorubicin-treated MEFs over NS MEFs controls (**Figure 4d, 4e**). γδ T cell killing was dose-dependent and time-dependent (**Supplementary Figure 5d, 5e, 5f**). Within 48 hours, mouse γδ T cells at 1:1 E:T eliminated approximately 80% of SEN MEFs (**Supplementary Figure 5e**). At 2:1 E:T, mouse γδ T cells killed the SEN population within approximately 40 hours (**Supplementary Figure 5f**). Our results show that murine γδ T cells target SEN MEFs. This suggests that our observations in human cells can be broadly generalized to mice.

**Figure 4.**
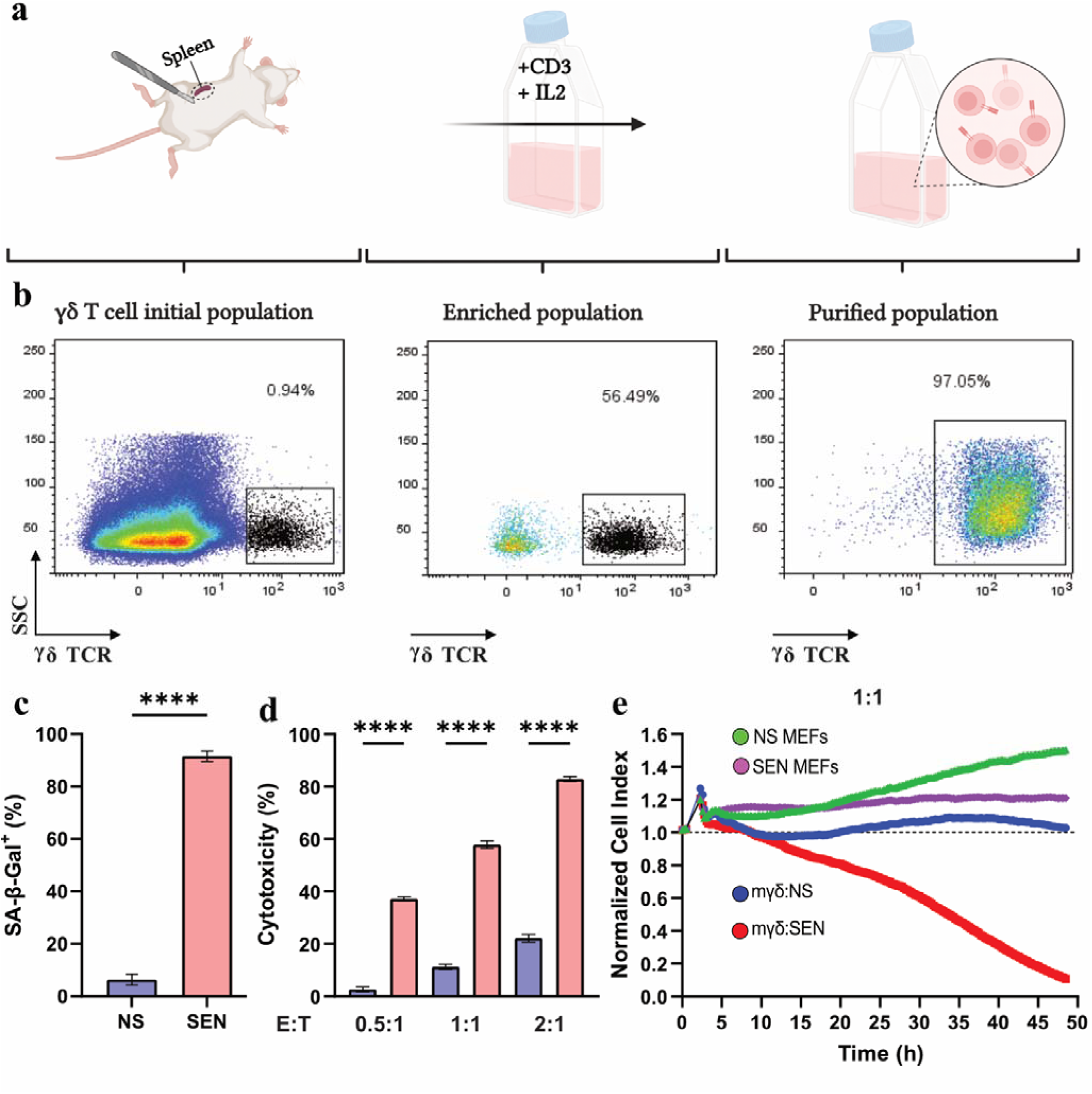
Mouse γδ T cells target senescent cells. (**a**) Schematic representation of the mouse γδ T cell enrichment strategy. Mouse γδ T cells were harvested from the spleen and immediately enriched using cell sorting. Mouse γδ T cells were expanded using IL-2 for 6-7 days. Mouse γδ T cells were purified using fluorescence-activated cell sorting and used for cytotoxicity assays. Created in https://BioRender.com. (**b**) Flow cytometry analysis of mouse γδ T cells isolated and enriched from the spleen. Left panel: Mouse γδ T cells identified within the splenocyte population before enrichment. Middle panel: Mouse γδ T cells population immediately after enrichment using fluorescence-activated cell sorting. Right panel: Highly purified mouse γδ T cell population obtained post-sorting and used for cell-killing assays. Representative data from experiments performed in at least three independent mouse spleen donors (n ≥ 3). (**c**) SA-β-Gal quantification in NS and doxorubicin-treated SEN MEFs. (**d**) Cytotoxicity of mouse γδ T cells to NS controls and doxorubicin-induced SEN at E:T ratios of 0.5:1, 1:1, and 2:1 after 24 hours. Representative data from experiments performed in at least three independent spleen donors (n ≥ 3); unpaired t-test. (**e**) Mouse γδ T cell 48-hour killing kinetics to NS controls and SEN at E:T ratio of 1:1. Representative data from experiments performed in at least three independent spleen donors (n ≥ 3).

**Figure 5.**
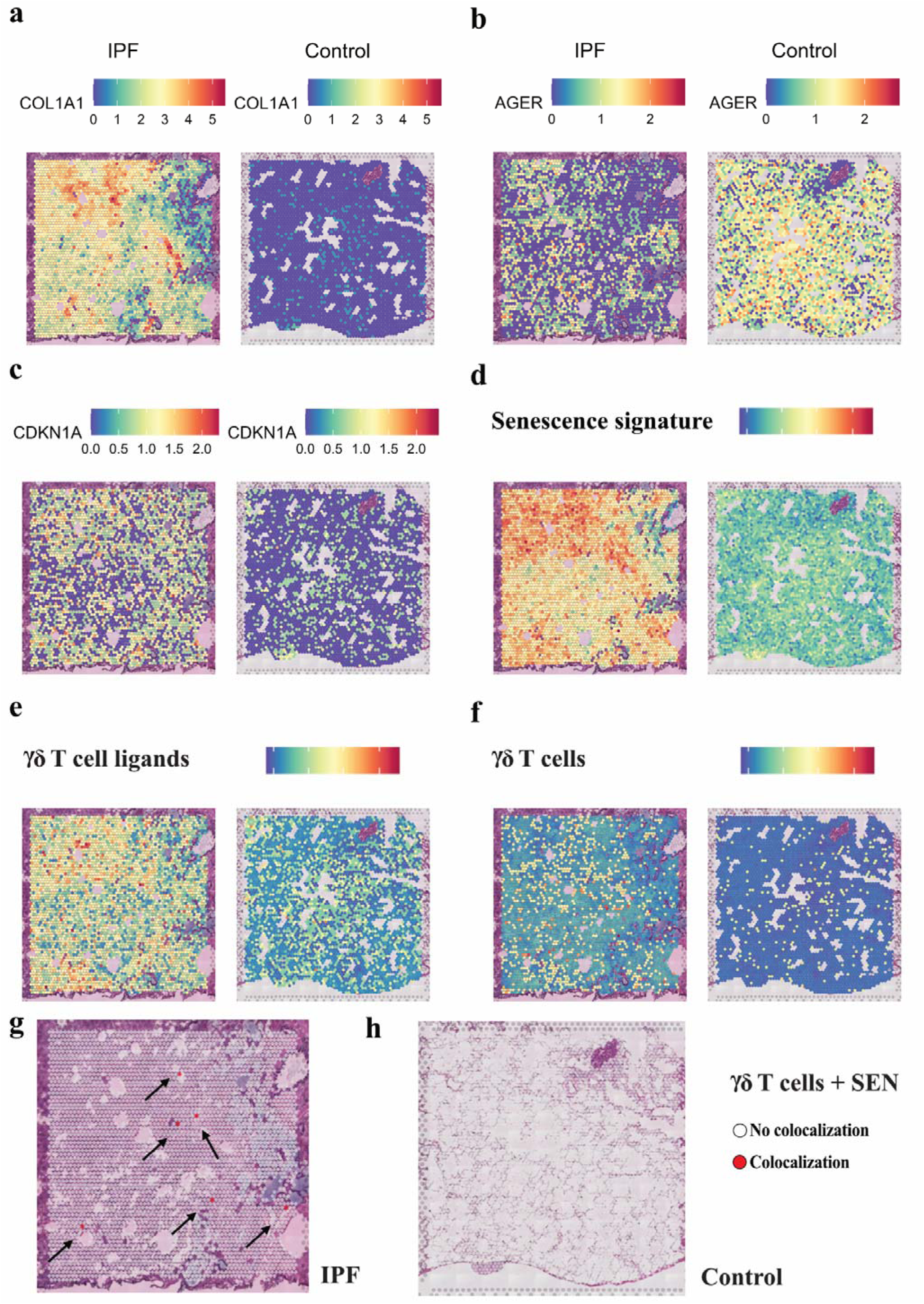
Human γδ T cell ligands colocalize with senescent cells and γδ T cells. (**a**) Validation of a pulmonary fibrosis gene signature in humans with IPF by spatial transcriptomics using COL1A1. Data was re-analyzed from Franzén et al., 2024 (PMID: 38951642). IPF = Freshly frozen human IPF lung tissue collected during transplantation with severe tissue remodeling. Control = Healthy control lung tissue (no known lung disease) collected postmortem. No tissue remodeling was observed in the healthy controls (PMID: 38951642). (**b**) Validation of a pulmonary fibrosis gene signature and loss of lung tissue homeostasis by spatial transcriptomics using AGER. (**c**) Validation of the presence of SEN (defined as CDKN1A-expressing cells) in IPF. (**d**) Senescence signature in the IPF lungs. A senescence signature was generated using the Saul Sen Mayo panel (PMID: 35974106). (**e**) γδ T cell ligand signature in the IPF lungs. A γδ T cell ligand signature was created using the genes ICAM1, MICA, BTN3A1, and PVR. (**f**) γδ T cell signature in the IPF lungs. A γδ T cell signature was created using the genes CD3E, TRDC, TRGC1, and TRGC2. (**g, h**) γδ T cell colocalization with SEN in the IPF lungs. γδ T cells (CD3E^+^ and TRDC^+^ spots) are colocalized with SEN (CDKN1A^+^ spots). (**g**) 6 spots with γδ T cells and SEN were identified in the IPF lungs; whereas (**h**) there were 0 spots with γδ T cells and SEN.

### γδ T cell ligands colocalize with senescent cells and **γδ** T cells in humans with idiopathic pulmonary fibrosis

Given these findings *in vitro,* we wanted to investigate further the physiological role of γδ T cells in humans. It is widely accepted that γδ T cells recognize and remove stressed cells (Deseke & Prinz, 2020). We hypothesized that if γδ T cells are engaged in senescent cell surveillance in humans *in vivo*, then γδ T cells would localize to SEN in human tissue that expresses ligands for γδ T cells. We analyzed a publicly available spatial transcriptomics dataset from human idiopathic pulmonary fibrosis (IPF) to explore this hypothesis (Franzen et al., 2024). IPF is a debilitating age-related lung disease characterized by progressive and irreversible fibrosis of the lung parenchyma, with a high burden of SEN of various types in affected tissue (Parimon et al., 2021). As expected, we observed a remarkable increase in spots enriched for COL1A1 in human IPF lung tissue relative to healthy controls, indicative of fibrosis (**Figure 5a**). Conversely, the IPF sample had a decrease in spots expressing AGER, a marker of lung homeostasis expressed by AT1 cells (Baek et al., 2021) (**Figure 5b**). The putative senescence marker CDKN1A/p21 was upregulated in IPF (**Figure 5c**), confirming the presence of senescent-like cells in the lungs of humans with IPF.

Consistent with our hypothesis, we observed that γδ T cell ligands were elevated in IPF samples compared to healthy control lung tissue (**Supplementary Figure 6a-f**). To achieve a higher level of resolution, we developed a senescence signature score using the “Saul Sen Mayo” panel (Saul et al., 2022) and identified lung regions with stronger senescence signatures (**Figure 5d**). These lung areas with a higher senescence signature also exhibited higher expression of γδ T cell ligands and p21 (**Figure 5c, 5e**), consistent with reports that SEN express stress-associated ligands (Yang et al., 2023). Finally, we created a γδ T cell signature and found that γδ T cells were close to SEN (**Figure 5f**). Given that each barcoded spot in spatial transcriptomics captures an average of one to ten cells, we examined whether SEN (CDKN1A^+^) colocalized with γδ T cells (CD3E^+^TRGC2^+^; the T cell receptor gamma constant). Further supporting our hypothesis, we observed that in IPF lungs, γδ T cells colocalized with SEN, whereas no such colocalization was observed in healthy matched control tissue (**Figure 5g, 5h**).

### γδ T cells expand during bleomycin-induced pulmonary fibrosis

Cellular senescence has been causally implicated in IPF’s pathogenesis, notably through the ability of senolytics therapies to ameliorate or reverse animal models of disease (Schafer et al., 2017). We used the well-established bleomycin-induced pulmonary fibrosis mouse model to investigate the role of γδ T cells *in vivo* (Liu et al., 2017; Walters & Kleeberger, 2008). Following bleomycin (BLM) instillation, mice exhibited weight loss, a proxy for successful BLM delivery and fibrotic lung injury (**Figure 6a**) (Schafer et al., 2017). Weight loss peaked around day 10 after BLM and stabilized afterward (**Figure 6a**). BLM-treated mice exhibited collagen deposition in the lungs, and a gene signature suggested pulmonary fibrosis compared to saline sham controls (**Supplementary Figure 7a, 7b**). Additionally, we observed a loss of spontaneous activity in the mice that received BLM, whereas sham mice remained active (**Supplementary Figure 7c, 7d**).

**Figure 6.**
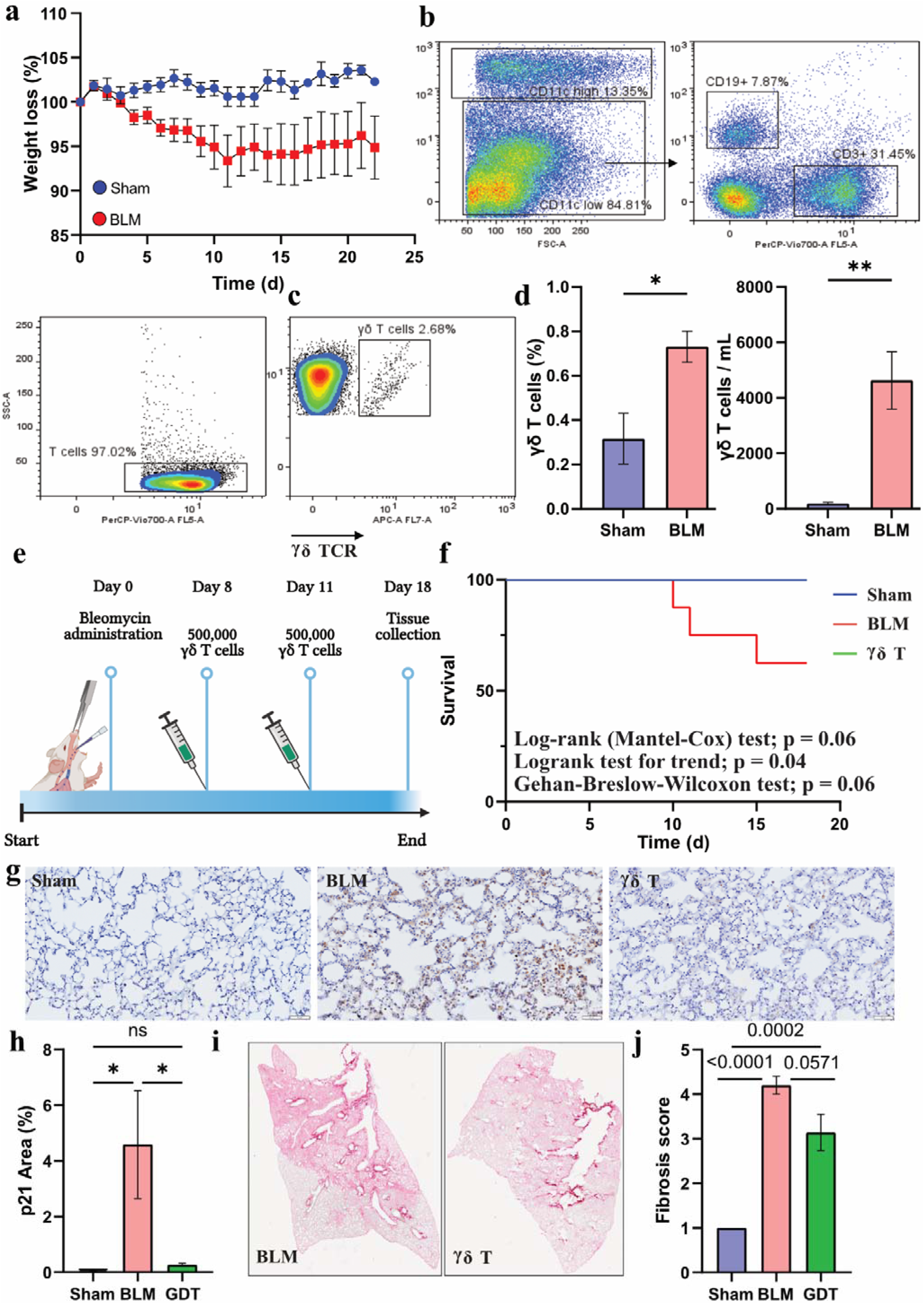
γδ T cells ameliorate pulmonary fibrosis and eliminate senescent cells. (**a**) Weight loss in BLM-treated mice. Male mice were administered BLM using oropharyngeal delivery. Weight was monitored daily; N = 5 sham-treated mice; N = 5 BLM-treated mice. (**b, c**) Representative images of the flow cytometry panel used to identify mouse γδ T cells after BLM insult. (**c**) Mouse γδ T cells were identified by gating them as Cd11c-low, CD3^+^, γδ TCR. (**d**) Mouse γδ T cells are recruited after fibrotic insult. The percentage and number of the total mouse γδ T cells was quantified by gating γδ TCR+ cells from the size and singlets gate in the BAL cells; N = 5 sham-treated mice; N = 5 BLM-treated mice. (**e**) Schematic representation of adoptive cell transfer of mouse γδ T cells. On day 0, mice were instilled 3 mg/kg of BLM using oropharyngeal delivery with a pipette. Sham-treated mice were instilled 50 μL of sterile saline. On days 8 and 11, a randomized subgroup of BLM-treated mice were injected 500,000 γδ T cells. Lungs were harvested 7 days after γδ T cell adoptive cell transfer. N = 6 sham-treated mice; N = 8 BLM-treated mice; N = 7 GFP^+^ γδ T cell transfer. (**f**) Kaplan-Meier survival probability after γδ T cell adoptive cell transfer. In the BLM-treated group, 3 out of 8 mice were excluded from the study. Out of 3 mice, 1 was found dead and 2 were euthanized due to humane endpoints. (**g**) Representative IHC p21 staining of the left lung. N = 6 sham-treated mice; N = 5 BLM-treated mice; N = 7 GFP^+^ γδ T cell transfer. Scale bar = 50 μm. (**h**) Quantification of p21^+^ area in the left lung. N = 5 BLM-treated mice; N = 7 GFP^+^ γδ T cell transfer. (**i**) Representative Picrosirius Red staining images of the left lung of BLM-treated and γδ T cell adoptive transfer. (**j**) Quantification of fibrosis in the γδ T cell adoptive transfer mice. Whole-lung Picrosirius Red slides were randomized and scored blindly on a scale of 1 to 5. 1 represents no fibrosis, whereas 5 represents a high degree of fibrotic lung area.

We designed a multi-color flow cytometry panel to interrogate immune populations post-BLM (**Figure 6b, 6c**). BLM-treated mice had increased total bronchoalveolar lavage (BAL) cells (**Supplementary Figure 7e**), reflecting increased inflammation, a hallmark of BLM-induced pulmonary fibrosis. We observed an increase in both the number and percent of T cells (CD3^+^) and γδ T cells (CD3^+^γδTCR^+^) compared to sham (**Figure 6d, Supplementary Figure 7f, 7g**). These findings align with studies reporting elevated γδ T cell numbers at multiple time points following BLM instillation (Pociask et al., 2011).

### γδ T cells attenuate bleomycin-induced pulmonary fibrosis and reduce p21-expressing cells

Senolytic therapies have been reported to ameliorate or reverse animal models of IPF (Schafer et al., 2017). Consistent with this and with our evidence that γδ T cells remove SEN, γδ T cell-deficient mice have been reported to exhibit severe BLM-induced lung injury and fibrosis (Pociask et al., 2011). We therefore investigated whether γδ T cell adoptive transfer in immunocompetent mice could ameliorate fibrosis and reduce senescence in the BLM-induced IPF mouse model. We isolated, enriched, and purified γδ T cells from the spleens of C57BL/6-Tg(UBC-GFP)30Scha/J (**Supplementary Figure 8a, 8b**). The GFP label allowed us to track the survival of γδ T cells in the fibrotic lungs. We induced IPF using BLM followed by adoptive cell transfer (ACT) of 500,000 γδ T cells on days 8 and 11 post-BLM instillation or vehicle (**Figure 6e**).

One of eight BLM-only mice died, and two were euthanized due to humane endpoints per IACUC protocol. All other mice that received ACT following BLM survived until all mice were euthanized one week after the final γδ T cell or vehicle administration (**Figure 6f**). BAL analysis confirmed the presence of GFP^+^ γδ T cells in all seven mice receiving ACT, with GFP^+^ cells staining positive for γδ TCR (**Supplementary Figure 8c, 8d**). Mice that received BLM plus ACT exhibited decreased lung inflammation relative to BLM-treated mice receiving vehicle instead of ACT (**Supplementary Figure 8e**). To assess whether γδ T cells had a senolytics effect, we stained for CDKN1A/p21 and observed a reduction in p21-expressing cells in the ACT group relative to mice that received BLM alone (**Figure 6g, 6h**). Finally, Picrosirius Red stain revealed that γδ T cells reduced the fibrosis score in mice receiving γδ T cells (**Figure 6i, 6j**). These results align with the report by Pociask and colleagues, supporting a protective effect of γδ T cells in BLM-induced lung fibrosis (Pociask et al., 2011). In summary, our findings reveal a novel role of γδ T cells in recognizing senescent cells (**Figure 7**).

**Figure 7.**
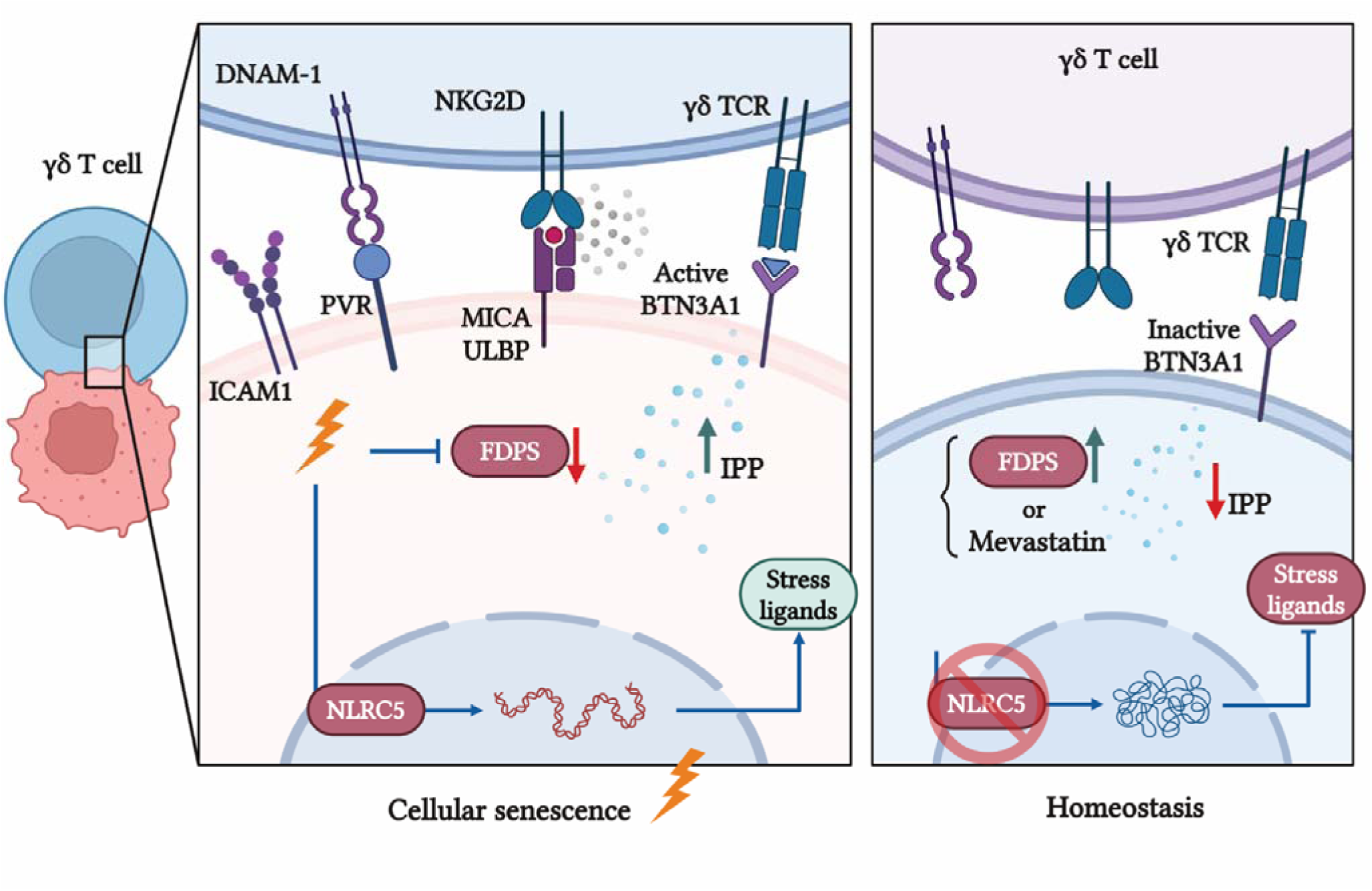
Proposed mechanism of γδ T cell recognition of senescent cells. Schematic of the proposed model of BTN3A1 and γδ T cell ligand presentation on the membrane of cells targeted by γδ T cells. Created in https://BioRender.com.

## Discussion

Senescent cells are characterized by a profound shift in their signaling and metabolic states, with increased stress signaling being a hallmark of their phenotype. This heightened stress signaling plays critical roles in their biology, interactions with the immune system, and contribution to age-related diseases. For instance, the persistent DNA damage caused by telomere shortening, oxidative stress, or genotoxic insults (e.g., chemotherapy or radiation) not only activates the DDR, resulting in the upregulation of cell cycle regulators such as p53, p21, and p16, but also increases the expression of MICA, MICB, ULBPs, and other MHC-I proteins (Marin, Boix, et al., 2023; Marin, Serrano, et al., 2023). These ligands can trigger their immune surveillance by NK and γδ T cells via NKG2D receptors (Deseke & Prinz, 2020).

The immune regulation of SEN by natural killer (NK) cells and macrophages was first demonstrated by Xue et al., showing that the reactivation of p53 induced senescence in tumor cells, which triggered immune-mediated clearance and tumor regression (Xue et al., 2007). The picture emerging in the immune regulation of senescence burden is highly complex, involving innate and adaptive immune leukocyte recognition and elimination of SEN, thereby maintaining tissue homeostasis and delaying the onset of diseases of aging (Arora et al., 2021; Binet et al., 2020; Hasegawa et al., 2023; Pereira et al., 2019; Sagiv et al., 2016). SEN play beneficial roles in tissue repair by promoting angiogenesis, stimulating tissue remodeling, and recruiting immune cells (de Magalhaes, 2024). The ability of immune cells to remove SEN after they have performed their regenerative functions is critical for ensuring tissue homeostasis and preventing excessive inflammation or fibrosis. However, several unknowns exist regarding the interplay between SEN and the immune system. For instance, the evolutionary drivers behind tissue-specific immune regulation, the reasons for the decline in SEN immunosurveillance with age, and how the antagonistic pleiotropy of senescence signaling balances beneficial and detrimental outcomes remain unclear. It is also poorly understood if and how different immune cells interact in an aging environment to eliminate the senescence burden. Thus, efforts are being made to develop a more comprehensive understanding of the immune-senescence axis. γδ T cells are a distinct subset of T lymphocytes that differ from conventional alpha beta (αβ) T cells in their T cell receptor (TCR) composition, function, and immune roles. They are a critical component of the immune system, operating at the intersection of innate and adaptive immunity. They play a significant role in combating infection, tumor immunity, and immune regulation (Deseke & Prinz, 2020). Given their unique stress-sensing capabilities, potent cytotoxic functions, tissue localization, and immunomodulatory plasticity, we investigated the immunoregulatory function of γδ T cells in surveilling SEN.

Unlike conventional αβ T cells, γδ T cells are uniquely equipped to detect and respond to stress-induced ligands on the targets through innate-like mechanisms (Deseke & Prinz, 2020). Our results show that in addition to NKG2D, γδ T cells engage with SEN using γδ TCR. This redundancy suggests a broader SEN immune surveillance capacity by this cell type relative to NK or αβ T cells. Moreover, our data shows that γδ T cells can eliminate SEN of diverse cell types of origin induced by various stressors. These features might enable flexible, context-dependent immune surveillance in different tissues.

To assess the SEN immune surveillance potential of γδ T cells, we isolated and expanded donor-derived Vγ9Vδ2 T cells, the predominant γδ T cell subset in peripheral blood, known for their ability to infiltrate tissues to remove target cells (Tosolini et al., 2017). However, we anticipate that Vδ1 T cells should also be capable of targeting SEN, as tissue-resident Vδ1 T cells are known to recognize “stress ligands” expressed on cells under stress conditions, such as transformed cancer cells or virus-infected cells, through receptors like NKG2D (Lawand et al., 2017).

Vγ9Vδ2 T cells express a range of chemokine receptors, such as CCR5, CCR2, CXCR3, and CXCR4, facilitating their migration toward inflamed or tumor-bearing tissues where corresponding chemokines are secreted (Glatzel et al., 2002; Prigione et al., 2004). SEN secrete several chemoattractants such as CCL5 and CXCL9, which recruit Vγ9Vδ2 T cells and other immune cells. This supports a physiological role for Vγ9Vδ2 T cells in the immune surveillance of SEN in aging or diseased tissues *in vivo*.

SEN are susceptible to ferroptosis, a form of cell death previously characterized in cancer cells. We recently demonstrated that SEN are primed to ferroptosis due to their avid accumulation of labile ferrous iron. This causes them to generate ROS through the Fenton reaction, producing oxidized lipids that initiate ferroptosis (Admasu et al., 2021). We further showed that SEN can be specifically targeted for destruction through this metabolic vulnerability (Admasu et al., 2021). These findings have suggestive implications for our present findings, as cell death via ferroptosis releases damage-associated molecular patterns (DAMPs) and lipid peroxidation products such as malondialdehyde (MDA) and 4-hydroxynonenal (4-HNE), secreted as SASP (Chen et al., 2023) signals that have been shown to activate γδ T cells (Schwacha et al., 2014).

Our data show the upregulation of MHC-I presentation machinery associated with NLRC5 and several BTN family genes. A previous study similarly reported *Nlrc5* to be upregulated in SEN MEFs (Marin, Boix, et al., 2023). Interestingly, a recent study using a genome-wide CRISPR KO approach identified NLRC5 as a key regulator of BTN3A1 expression (Mamedov et al., 2023). γδ T cells also rely on the accumulation of IPP and the GTPase RhoB to effect their cytotoxic potential through BTN3A1 (Gu et al., 2017; Sebestyen et al., 2016). Pharmacological inhibition of FDPS, a mevalonate pathway enzyme responsible for converting the phosphoantigen IPP into farnesyl diphosphate (FPP), leads to the intracellular accumulation of IPP and is sufficient to activate Vγ9Vδ2 T cells *in vitro* and in murine xenograft cancer models *in vivo* (Liou et al., 2022). Similarly, FPDS KO cells trigger high γδ T cell recognition, and ZOL inhibition of FDPS leads to an increase in BTN3A1 membrane localization (Mamedov et al., 2023). We show that SEN have lower levels of FDPS compared to NS controls. Additionally, in our study, inhibition of the mevalonate pathway with mevastatin (and thus lower levels of intracellular IPP) was sufficient to reduce BTN3A1 expression and selective seno-toxicity by γδ T cells.

In a previous study, we reported that SEN utilize GPX4 to lower oxidative stress and resist their intrinsic vulnerability to ferroptotic death (Admasu et al., 2021). However, GPX4 synthesis depends on the IPP produced by the mevalonate pathway (Weaver & Skouta, 2022). We hypothesize that SEN’s higher metabolic demands for GPX4 may lead to increased production of IPP and other phosphoantigens, which upregulate the expression of CD277, an activating ligand for γδ T cells (Harly et al., 2012). This potential mechanistic connection between ferroptosis resistance and enhanced immune-mediated clearance of SEN opens up novel therapeutic avenues to target SEN, which remain to be investigated. Additionally, other potential mechanisms of IPP accumulation in SEN beyond the mevalonate pathway merit investigation, as multiple relevant metabolic pathways are deregulated in SEN (e.g., an increase in cholesterol efflux and uptake mediated by CD36). This is an intriguing possibility and must be investigated in future studies.

Cancer cells are known to secrete IPP and activate γδ T cell chemotaxis (Ashihara et al., 2015). This suggests the possibility that SEN similarly release IPP to the extracellular space, further priming γδ T activation and killing. Given that ABCA1 is a key enzyme for the release of IPP and activation of γδ T cells in dendritic cells (Castella et al., 2017) and that ABCA1 is upregulated in SEN (Roh et al., 2023), it is plausible that SEN not only presents γδ T cell ligands but also activates these cells through the release of IPP into the extracellular space.

According to an established protocol, our study used the nitrogen-containing bisphosphonate drug zoledronate to enrich γδ T cells with an effector memory phenotype (Kondo et al., 2008). The ability of ZOL to expand and activate γδ T cells is grounded in its inhibition of FDPS, leading to the accumulation of its substrate phosphoantigen IPP (Thompson & Rogers, 2004). γδ T cell expansion and activation have also been observed in human cancer patients treated with ZOL (Dieli et al., 2003; Naoe et al., 2010). Consistent with an effect on SEN, ZOL reduced circulating SASP factors and improved grip strength in aged mice, accompanied by decreased protein levels of p16, p21, and SASP factors in their preosteoclasts (Samakkarnthai et al., 2023). The investigators provided some evidence for a direct senolytic effect of ZOL.

Still, we speculate that the *in vivo* results may also have resulted from γδ T cell activation in treated mice. Moreover, secondary analysis of the randomized controlled HORIZON trial of ZOL for fracture prevention and pharmacoepidemiological studies has reported unexpected benefits of ZOL treatment, including increased survival (Center et al., 2020). These findings suggest that ZOL may expand and activate γδ T cells *in vivo*, enhancing SEN immunosurveillance and improving survival.

Moreover, several groups have reported that the frequency and absolute number of γδ T cells in the periphery decline with age (Kurioka & Klenerman, 2023). Based on our findings and those of others just reviewed, it is plausible that this age-related decline may partly be responsible for the accumulation of SEN in aging tissues.

Similarly, administration of the CD1d-binding glycolipid antigen α-galactosylceramide (α-GalCer) has been shown to result in ablation of SEN in high-fat fed obese mice and the BLM mouse model of idiopathic pulmonary fibrosis, leading to improved functional outcomes and, in the latter case, survivorship (Arora et al., 2021). α-GalCer is known to activate invariant natural killer cells (iNKT cells) *in vivo* in mice and humans, and these investigators provided evidence that activated iNKT cells were engaged in immune surveillance in these mouse models and human iNKT cells in culture (Arora et al., 2021). However, these results may additionally implicate γδ T cell immunosurveillance, as the complete immunological response to α-GalCer in a murine cancer model was shown to depend on a bystander effect on γδ T cells, which increased their cytotoxic activity against tumor cells after α-GalCer injection downstream of the activation of iNKT (Paget et al., 2012). This again suggests that part of the observed SEN immunosurveillance attributed to iNKT may lie with γδ T cells (Paget et al., 2012). Our findings and previous reports with ZOL and α-GalCer suggest that the γδ T cell senescent cell surveillance reported here may be operative *in vivo* and therapeutically tractable.

In summary, this study provides the basis for investigating γδ T cells in the context of aging, age-related diseases, and senescence-driven pathologies. We present an optimized protocol for isolating, activating, enriching, and purifying human and mouse γδ T cells. Additionally, we share a resource to understand human lung SEN through snRNA-Seq. Future investigations must elucidate the role of senescence stress-associated ligands in γδ T cell surveillance of senescent cells and how to exploit this underinvestigated immune population to improve human health and lifespan.

## Methods

### Cell culture

IMR-90 fibroblasts (ATCC, USA; Cat# CCL-186) were maintained at 37°C in humidified air containing 5% CO_2_ and 3% O_2_. Fibroblasts were used at population doubling level (PDL) 25-63 and maintained in complete media containing Dulbecco’s Modified Eagle’s Medium (DMEM) (Corning; Cat# 10-013-CV) supplemented with 10% Fetal Bovine Serum (FBS) (Millipore Sigma, USA; Cat# F4135) and 1X Penicillin– Streptomycin (Corning; Cat# 30-001-CI). Mycoplasma contamination was routinely tested using a standard PCR test. Cumulative PDL was calculated using the following equation:

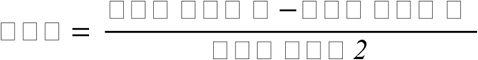

where H is the number of cells at harvest and S is the number of cells seeded.

Mouse Embryonic Fibroblasts (MEFs) were isolated from the mouse embryo as described elsewhere (Samakkarnthai et al., 2023). Shortly, embryos were finely chopped at embryonic day 12 to 13 and cultivated in supplemented DMEM medium. MEFs were used at passage (P) P2 to P8 and kept at 37°C in humidified air containing 5% CO_2_ and 3% O_2_. Mycoplasma contamination was routinely tested using a standard PCR.

### Senescence induction

Human IMR-90 fibroblasts were treated with 300 nM of doxorubicin hydrochloride (Millipore Sigma, USA; Cat# 504042) in DMEM complete media for 24 hours and maintained in culture. For an alternative mode of senescence induction, IMR-90 fibroblasts were treated with 5 μM etoposide (ApexBio; Cat# A1971) for 48 hours in DMEM complete media and maintained in culture. MEFs were treated with 150 nM of doxorubicin hydrochloride (Millipore Sigma, USA; Cat# 504042) in DMEM complete media for 24 hours and maintained in culture. HUVECs were treated with 100 nM of doxorubicin hydrochloride (Millipore Sigma; USA; Cat# 504042) in DMEM complete media for 24 hours and maintained in culture. SEN were used for experiments 7-10 days after doxorubicin or etoposide treatment. Replicative senescence was confirmed by complete cessation of cell proliferation, clear morphological changes, and high (>80%) SA-β Gal^+^ cells, generally around PDL 58-63.

### Senescence-associated **β**-galactosidase activity

SA-β Gal activity was detected as described elsewhere (Dimri et al., 1995) using a commercial kit (Senescence Detection Kit; Cat# ab65351) and following the providers’ protocol.

### Immunofluorescence

Cells were plated at a single-cell density prior to senescence induction.10 days after senescence induction, cells were fixed with 4% PFA for 15 minutes at room temperature (RT). For intracellular staining, cells were then permeabilized in 0.2% Triton X-100 for 15 minutes and blocked in 5% goat serum (GS) in PBS for 1 hour. Primary antibodies were added and left overnight at 4°C on a shaker. Primary antibodies used were γH2AX (Novus; P Ser139 3F2; Cat# NB100-74435) (1:1000 dilution), anti-HMGB1 antibody (Abcam; Cat# ab18256) (1:1000 dilution), and RhoB Recombinat Polyclocal (Invitrogen; Cat# 711274) (1:500 dilution). CD277 immunofluorescence was performed on non-permeabilized cells using BTN3A1 Polyclonal antibody (Proteintech, Cat# 25221-1-AP) (1:1000 dilution). After incubation with primary antibodies, cells were incubated with fluorescent secondary antibodies and Hoechst (Invitrogen; Cat# H3570) (1: 2000 dilution) for 1 hour at RT in the dark. Secondary antibodies used were Goat anti-Rabbit IgG (H + L) Cross-Adsorbed Secondary Antibody, Alexa Fluor 546 (Invitrogen; Cat# A-11010) and Goat anti-Mouse IgG (H+L) Highly Cross-Adsorbed Secondary Antibody, Alexa Fluor™ 488 (Invitrogen; Cat# A-11029). Images were acquired with an EVOS cell imaging system at 20x and 40x magnification. ImageJ software was used to measure fluorescent intensity per cell.

### Human **γδ** T cell isolation and enrichment

γδ T cells were isolated and enriched from human blood using a previously published protocol (Kondo et al., 2008). The blood samples were obtained from donors in heparin-coated vacutainers. All subjects provided informed written consent. Blood samples were diluted with sterile 1X PBS (Corning; Cat# 21-031-CV) and combined with Lymphocyte Separation Medium (Corning; Cat# 25-072-CI) followed by density gradient centrifugation, according to the manufacturer’s instructions. An aliquot of PBMC was analyzed for the proportion of γδ T cells by flow cytometry. The remainder of PBMCs were cultured in RPMI complete media, 10% Fetal Bovine Serum (FBS) (Millipore Sigma, USA; Cat# F4135) with 500 IU/ml human rIL-2 (recombinant Interleukin-2) (TECIN teceleukin; Cat# Bulk Ro 23-6019) and 5uM zoledronic acid (ZOL) (Tocris; USA Cat #6111) for 24 hours. The cells were expanded with rIL-2 for 10 to 14 days with fresh media changed every other day, before coculturing with senescent or non-senescent IMR-90 fibroblasts. After 10 - 14 days of expansion, γδ T cells were negatively selected using γδ T cells Isolation Kit (Miltenyi Biotec; Cat# 130-092-892) or positively selected using MACSQuant® Tyto® Cell Sorter.

### Murine **γδ** T cell isolation and enrichment

All experiments involving mice were approved by the Institutional Animal Care & Use Committee (IACUC). Mouse γδ T cells were isolated and expanded from the spleen of multiple mice using a modified version of a protocol described elsewhere (Williams et al., 2022). Briefly, 6 to 12-month-old mice were sacrificed using a CO_2_ chamber, followed by cervical dislocation. Spleens were collected and transferred to a sterile 50 mL Falcon tube containing cold PEB Buffer (PBS, 0.5% bovine serum albumin (BSA), and 2 mM EDTA) and washed 3 times. Spleens were transferred into a gentleMACS C Tube (Miltenyi Biotec; Cat# 130-093-237) containing 5 mL of cold PEB Buffer. Spleens were mechanically dissociated for one minute using the gentleMACS Octo Dissociator with Heaters (Miltenyi Biotec; Cat# 130-096-427). Dissociated spleens were run through a cell strainer to remove remaining pieces of tissue (Miltenyi Biotec; Cat# 130-101-812), and gentleMACS C Tubes were washed thoroughly with PEB buffer. Single-cell splenocytes were pelleted by centrifugation at 300g × 10 minutes, and red blood cells were lysed using sterile RBC Lysis Buffer (Zymo Research; Cat# R1022-2-100). Then, splenocytes were labeled for positive sorting using MACSQuant® Tyto® Cell Sorter using TCRγ/δ antibody, anti-mouse, APC, REAfinity™ (Miltenyi Biotec; Cat# 130-123-851). Enriched γδ T cells were cultured for 24 hours in OKT Anti-Mouse CD3-coated wells (Tonbo Biosciences; Cat# 40-0032-U100) and 1000 U/mL rIL-2 (recombinant Interleukin-2) (TECIN teceleukin; Cat# Bulk Ro 23-6019) in RPMI-1640 Medium (ATCC; Cat# 30-2001), supplemented with 10% FBS (Millipore Sigma, USA; Cat# F4135), 1X Penicillin–Streptomycin (Corning; Cat# 30-001-CI), and 55mM β-Mercaptoethanol (Gibco; Cat# 21985023). Then, for 48 hours with OKT Anti-Mouse CD3-coated wells and 100 U/mL rIL-2. Finally, for five more days with 100 U/mL rIL-2. γδ T cell purity was assessed on day 6 and cells were positively sorted on day 6-7 to achieve <90% purity using the MACSQuant® Tyto® Cell Sorter.

### Flow cytometry

Adherent cells were detached using trypsin. Cells were resuspended in 100 μl of PBS. Cells were then incubated with APC-conjugated anti-human CD3 antibody (Miltenyi Biotec; Cat# 130-113-135), CD3 Antibody, anti-human, VioBlue, REAfinity (Miltenyi Biotec; Cat# 130-114-519), TCR α/β Antibody, anti-human, APC, REAfinity (Miltenyi Biotec; Cat# 130-113-535), FITC-conjugated anti-human γδ TCR antibody (Miltenyi Biotec; Cat# 130-113-503), TCR γ/δ Antibody, anti-human, PE, REAfinity (Miltenyi Biotec; Cat# 130-113-512), PE-conjugated TCR Vδ1 Antibody (Miltenyi Biotec; Cat# 130-120-440), FITC-conjugated TCR Vδ2 Antibody (Miltenyi Biotec; Cat# 130-111-009), CD56 Antibody, anti-human, Vio Bright R667, REAfinity (Miltenyi Biotec; Cat# 130-114-553), FITC-conjugated CD56 antibody (Miltenyi Biotec; Cat#130-113-312), and APC-conjugated CD56 antibody (Miltenyi Biotec; Cat#130-113-312) strictly following the provider instructions on ice for 30 minutes. Cells were washed with 1 ml of ice-cold PBS and resuspended in 200 μl of ice-cold PBS and analyzed on the MACSQuant10, Miltenyi Biotec flow cytometer. Cell viability was determined by 7AAD (Miltenyi Biotech; Cat# 130-111-568) staining, and live cells were gated for downstream analysis. Data were analyzed using Flowlogic software (Miltenyi Biotech; Germany).

### Real-time cytotoxicity assay (xCELLigence)

Cytotoxicity toward SEN was measured using the xCELLigence platform which uses a cellular impedance-based measurement as a proxy of cell attachment and cell number. Positive and negative controls were used to calculate viability and cytotoxicity. Immune cells in co-culture remain in suspension and do not adhere to the wells, and as such contribute negligibly to the impedance reading. Initially, 50μL medium was added to E-Plates 96 (Agilent; Cat# 300-600-910) for measurement of background values. SEN and their respective NS controls were seeded in an additional 100μL medium at a density of 10,000 cells per well. Cell attachment was monitored using the RTCA MP (Agilent) instrument and the RTCA software (Agilent) until the plateau phase was reached, typically after 24 hours. γδ T cells were added on top of the SEN and NS at different E:T ratios. Cells treated with 0.2% Triton X-100 were used as a 100% dead cell positive control for the cytotoxicity assay. Upon addition of γδ T cells, impedance measurements were performed every 15 min for up to 48 hours. All experiments were performed with at least three independent replicates for each donor and cell type. Changes in impedance were expressed as a cell index (CI) value, which derives from relative impedance changes corresponding to cellular coverage of the electrode sensors, normalized to baseline impedance values with medium only. Percent cytotoxicity was determined based on the relative CI value. To analyze the acquired data, CI values were exported, and the percentage of lysis was calculated in relation to the control cells lacking any γδ T cells.

### Animal care

All protocols with mice were approved by the Institutional Animal Care & Use Committee (IACUC). Mice were given food ad libitum and kept under a 12-hour light/dark cycle. 4-6-month-old C57/B6 mice were purchased from The Jackson Laboratory. For modelling idiopathic pulmonary fibrosis, only male C57BL/6J were used. For GFP^+^ γδ T cells, male and female C57BL/6-Tg(UBC-GFP)30Scha/J were purchased from The Jackson Laboratory and bred in house. All purchased mice were kept in quarantine for two weeks. All mice were checked daily by a trained specialist.

### Open field

Loss of spontaneous activity after BLM instillation was assessed using open field (Noldus EthoVision). Mice were acclimated to the testing room for at least one hour. Mice were placed in the EthoVision arena for 10 minutes and allowed to freely explore and move. Spontaneous activity was recorded and analyzed using the Noldus Ethovision software. Total distance, in centimeters, was measured by tracking the center-point of each animal and its movement in the arena.

### Bleomycin-induced pulmonary fibrosis *in vivo* mouse model

Pulmonary fibrosis was induced using the oropharyngeal tongue-pulling technique, as described elsewhere (Liu et al., 2017; Walters & Kleeberger, 2008). Bleomycin sulfate (MedChem Express; HY-17565, powder 5mg) was freshly prepared in saline for each instillation. On day 0, isofluorane-anesthesized male mice received 2.5 mg/kg or 3 mg/kg (final volume 50 μL) of Bleomycin sulfate. Following BLM delivery, mice were weighed and monitored daily for signs of deterioration. Mice were given gels and additional food if mild or moderate BLM-induced weight loss was observed. If severe weight loss or signs of distress and pain were observed mice were excluded from the study and humanely euthanized. Mice that lost less than 10% of weight by day 8 after BLM instillation were excluded from the study. 500,000 γδ T cells were administered on days 8 and 11 using the same oropharyngeal tongue-pulling technique. A dose of 100,000 IU of IL-2 was administered intraperitoneally (i.p.) during the ACT to all BLM-receiving mice. No negative effects on weight or behavior were observed from the γδ T cell ACT. After 18 or 21 days, mice were euthanized, and lung tissue was collected. The left lungs were collected for histology whereas the right lungs were immediately flash frozen for RNA isolation and gene expression.

### Murine Bronchoalveolar Lavage

Bronchoalveolar lavage (BAL) was done following a previously described protocol (Van Hoecke et al., 2017). After mouse euthanasia, BAL was collected by using 2 x 0.8 mL ice-cold PBS buffer. BAL fluid was frozen at -80°C, and cells were pelleted and stained for flow cytometry. Cells were stained for 30 minutes at 4°C, washed, and analyzed using flow cytometry as described above (MACSQuant10, Miltenyi Biotec). The following antibodies were used, CD3ε Antibody, anti-mouse, PerCP-Vio 700 (Miltenyi Biotec; Cat# 130-120-174), CD19 Antibody, anti-mouse, PE-Vio 770 (Miltenyi Biotec; Cat# 130-123-881), CD11c Antibody, anti-mouse, VioBlue (Miltenyi Biotec; Cat# 130-102-413).

### Histology

The left lung was collected and fixed using 10% neutral buffered formalin (Sigma Aldrich; Cat# HT501128) for 24 hours. Lungs were washed one time in PBS and kept in ethanol 70% prior to paraffin-embedding. Formalin-fixed paraffin-embedded (FFPE) tissue blocks were prepared by Zyagen (CA, USA). For Masson-trichrome staining (VitroVivo Biotech; VB-3016), 5 μm sections were prepared, deparaffinized, and stained strictly following the manufacturer’s instructions. For Picrosirius Red staining, 5 μm sections were prepared, deparaffinized, and rehydrated. Sections were treated with phosphomolybdic acid aqueous solution 0.2% for 5 minutes and washed by a 5-minute tap water rinse. Then, 0.1% saturated picric acid was used to stain the sections for 2 hours at RT. Finally, the sections were washed in freshly prepared 0.015N hydrochloric acid for 3 minutes, rinsed in 70% ethanol, rehydrated, and mounted (Fisher Chemical Permount Mounting Medium; Cat# SP15-100). Sections were imaged using the Olympus SlideView VS200. Picrosirius Red-stained sections were randomized and the fibrosis score on scale 1-5 was determined by a blinded researcher.

### Immunohistochemistry

5 μm FFPE sections were used for p21 IHC staining to quantify senescence burden. FFPE sections were deparaffinized and re-hydrated. Antigen retrieval was performed using boiling-temperature Tris-HCl (G Biosciences; Cat# RC-105) buffer at pH 10 for 10 minutes. Endogenous peroxidases were inactivated with freshly prepared 3% hydrogen peroxide (Sigma-Aldrich; Cat# 216763) for 10 minutes. Sections were blocked for one hour at RT using 5% GS (Gibco; Cat# 16210-064) in TBST (Bio-Rad; Cat# 1706435) and stained using 1:50 p21 (Thermo Fisher Scientific; Cat# 14-6715-81) overnight at 4°C. Sections were washed and incubated with HRP-conjugated goat anti-rabbit secondary antibodies (ImmPRESS HRP goat anti-rabbit HRP IgG polymer kit, Vector Laboratories; Cat# MP-7451-15). DAB substrate was used to visualize p21 stain (ImmPACT DAB substrate peroxidase kit, Vector Laboratories; Cat# SK-4105). Counterstaining with 1:5 hematoxylin was done for 1.5 minutes (Vector Laboratories; Cat# H-3401; Hematoxylin and Eosin Stain Kit; Cat# H-3502). Sections were washed, rehydrated, mounted (Fisher Chemical Permount Mounting Medium; Cat# SP15-100) and imaged using the Olympus SlideView VS200. Four random fields of view were taken and evaluated for p21 staining using ImageJ.

### RNA isolation from lungs

After dissection, lungs were flash-frozen using liquid nitrogen and kept at -80C. Lungs were homogenized into a fine powder using cryogenic grinding with a pestle and a mortar and liquid nitrogen. Powdered lungs were kept in RLT Buffer supplemented with β-Mercaptoethanol, and further homogenized using a 21-Gauge Syringe (BD; Cat# 305165). RNA isolation was done following the manufacturer’s instructions (QIAGEN, RNeasy Plus Mini Kit; Cat# 74136; QIAshredder; Cat# 79656). All samples were tested for RNA concentration and purity with Nanodrop (Thermo Fischer), and one BLM sample was excluded due to poor RNA yield. qPCR was done using Sybr Dye (AzuraQuant Green Fast qPCR Mix LoRox, Cat# AZQ-Lo-2000).

### Quantitative PCR

RNA from NS and SEN cells was isolated following the manufacturer’s protocol (Zymo Research, Quick-RNA Miniprep Kit; Cat# R1055). The AzuraQuant cDNA Synthesis Kit (Azura Genomics; Cat# AZ-1996) was used to prepare cDNA, and qPCRs were done using Sybr Dye (AzuraQuant Green Fast qPCR Mix LoRox; Cat# AZQ-Lo-2000).

Human genes:

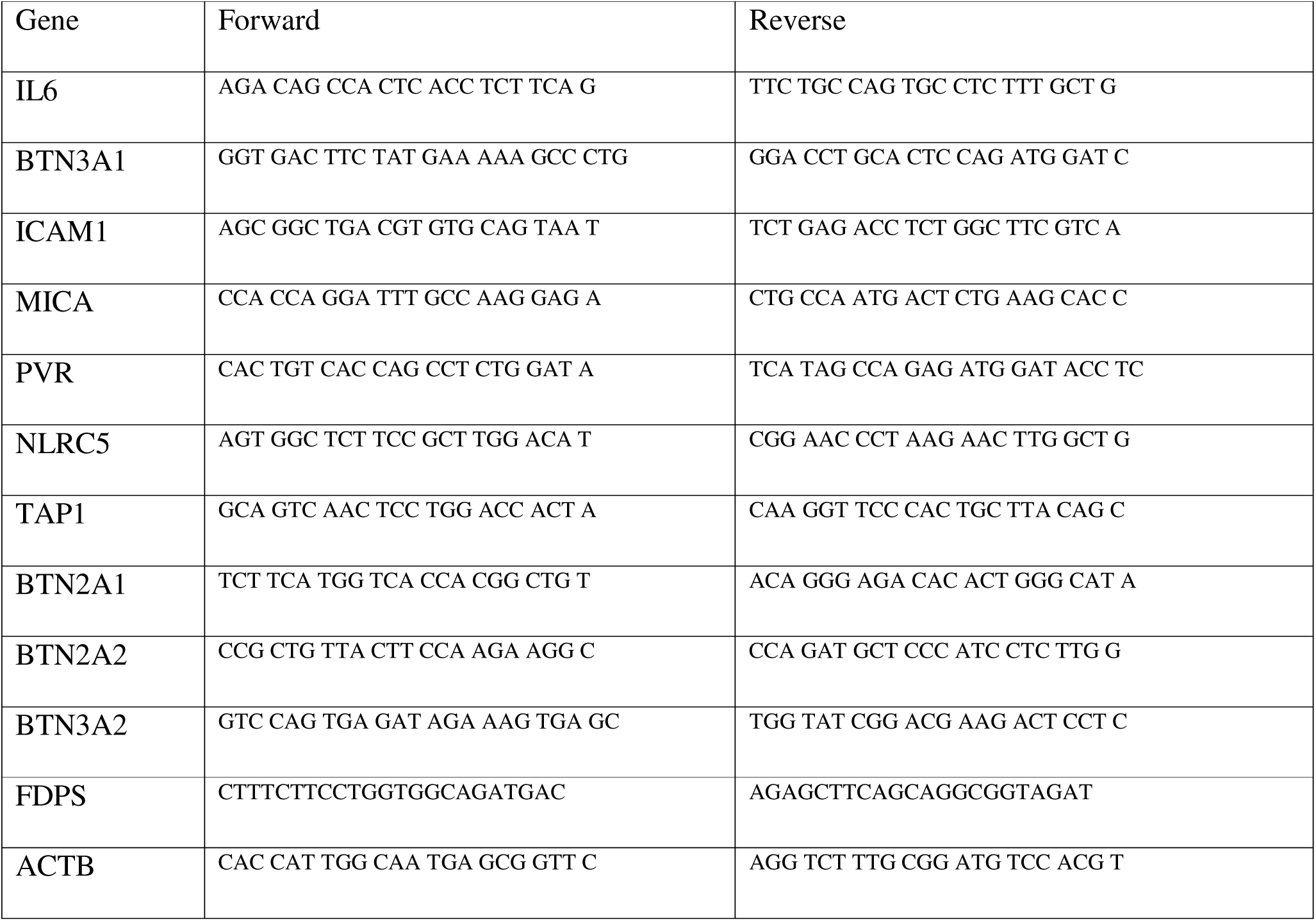

Mouse genes:

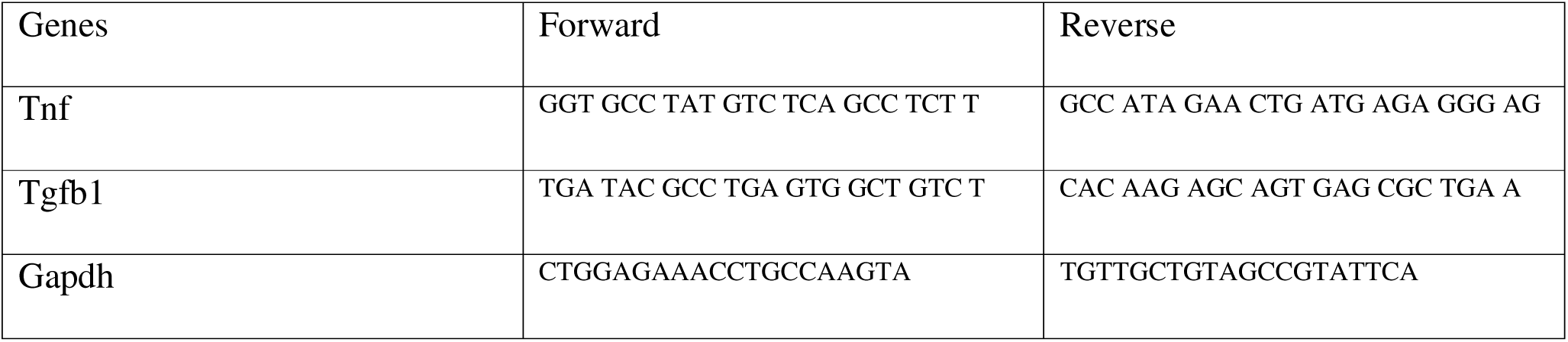

### Single-nucleus RNA Sequencing, quality control, and analysis

Human fibroblast samples were transported on dry ice from the SENS Research Foundation to the Buck Institute for Research on Aging and stored at –80°C upon arrival. For nuclei isolation, fibroblasts were transferred into pre-chilled 1.5 mL Eppendorf tubes and dissociated using the 10x Genomics Nuclei Isolation Kit (10X Genomics; Cat# PN: 1000494). Lysis was performed for 3 minutes, followed by two resuspension washes, according to the manufacturer’s instructions. Isolated nuclei were fixed at 4°C for 24 hours in 4% paraformaldehyde (PFA), prepared by diluting 37% PFA (Fisher BioReagents; Cat# BP531-25) in nuclease-free water. Nuclei concentration was determined using the Countess II Automated Cell Counter (Thermo Fisher Scientific, Waltham, MA, USA) with propidium iodide staining (Invitrogen, Waltham, MA, USA). Single-cell libraries were prepared using the Chromium Next GEM Single Cell Fixed RNA Sample Preparation Kit (10X Genomics; Cat# PN: 1000323) on the Chromium X instrument, following the manufacturer’s protocol. Library quality and fragment size distribution were assessed using the Agilent TapeStation (Agilent Technologies, Santa Clara, CA, USA). Sequencing was performed by SeqMatic (Fremont, CA, USA) on one lane of a NovaSeq X 10B system using 2 × 150 bp paired-end reads. Barcodes, matrix, and feature files were generated using Cell Ranger. The snRNA-Seq. was processed, explored, and visualized using Trailmaker/Cellenics® community instance (https://scp.biomage.net/) hosted by Biomage (https://biomage.net/). For ambient RNA discrimination, a False Discovery Rate (FDR) was calculated for each cell barcode using the emptyDrops function. Barcodes with an FDR value < 0.1 were used. Additionally, a hard Unique Molecular Identifier (UMI) threshold of > 500 UMIs allowed further distinction between real cells and empty droplets. Barcodes with over 10% mitochondrial gene content were filtered out. Outliers in the relationship between the number of expressed genes and the number of UMIs were identified and excluded by fitting a linear regression model and using a stringent 0.001 p-value criteria. Barcodes with a high probability of being a doublet were excluded using scDblFinder. After quality control, 668 non-senescent cells and 626 SEN were analyzed. To correct for batch effects, the Harmony integration method was selected. Data was LogNormalized, and the 2000 top highly variable genes (HVGs) were selected. For two-dimensional data visualization, the top 20 principal components were used and data was embedded using Uniform Manifold Approximation and Projection (UMAP). Cell clusters were identified using the Louvain algorithm with a resolution of 0.3.

### Data representation and statistical analysis

Statistical analysis was conducted using Graph Pad Prism 10. Images were analyzed using ImageJ. Flow cytometry experiments were analyzed using FlowLogic. All data are presented as means ± SEM, unless specified in the figure legend. All experiments have been performed at least 3 independent times. Comparisons between groups were performed using unpaired t-test, 2-tailed Student’s t test, 1- or two-way ANOVA, as appropriate with appropriate correction for multiple comparisons. Statistical parameters can be found in figure legends.

## Acknowledgements

We acknowledge Esmeralda Jimenez (E.J.) for her expertise in animal care. E.J. conducted daily mouse checks and maintenance, provided insights on animal welfare, and advised on experimental procedures. We acknowledge Prof. Simon Melov (Buck Institute, California, USA) for allowing us to use equipment in his lab.

## Data availability

Data will be made available upon reasonable request. The spatial transcriptomics data can be found at https://www.ebi.ac.uk/biostudies/studies/S-BSST1410. Additionally, the single-cell RNA Sequencing data from WI-38 SEN was found at GSE226225.

## Author contributions

G.M.L., T.A., and A. Sharma wrote the manuscript. M.R. edited the manuscript. G.M.L. and T.A. performed most experiments and analyzed results. A.B., A. Shankar, I.C., performed experiments and analyzed results. T.T. and A.B. prepared samples for sequencing. A. Sharma, T.A., and G.M.L. designed experiments. A. Sharma supervised the project. A. Sharma obtained funding. All authors have read and edited the manuscript. All authors contributed to the discussion. All authors agree with the final version of the manuscript.

