## Supplementary material for "γδ T Cells Target and Ablate Senescent Cells in Aging and Alleviate Pulmonary Fibrosis": Supplimental data

**Gamma Delta (γδ) T Cells Target Senescent Cells**

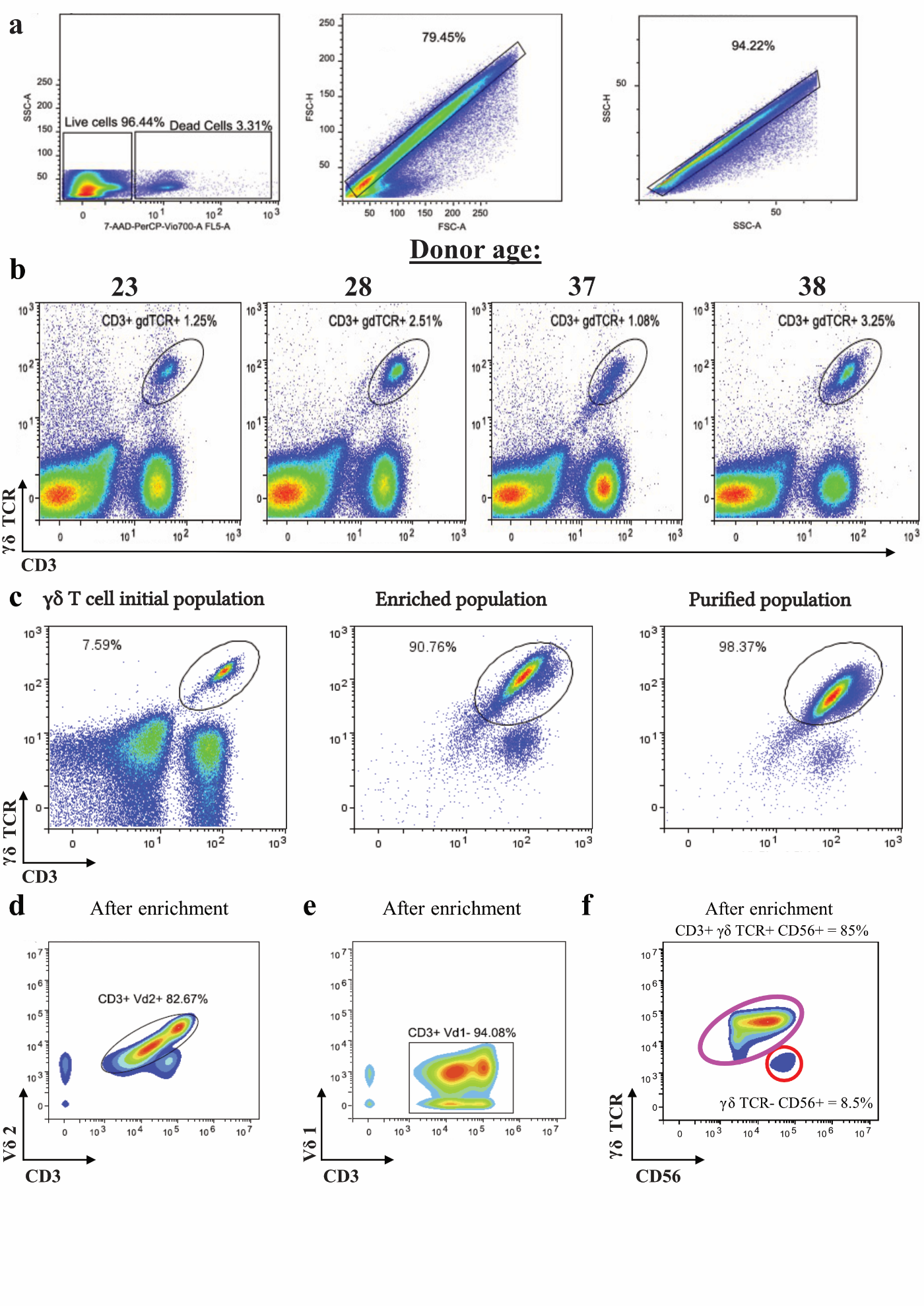

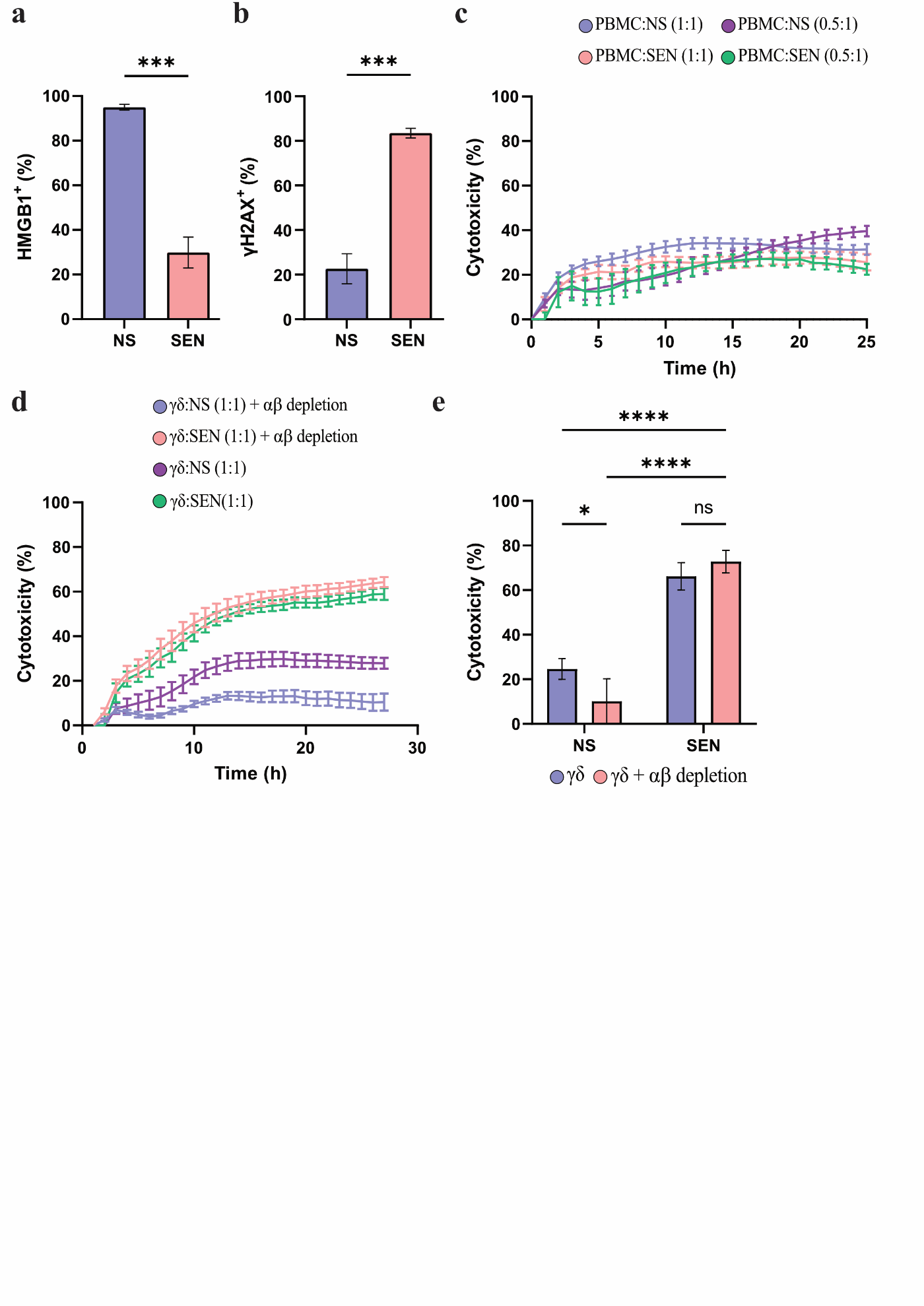

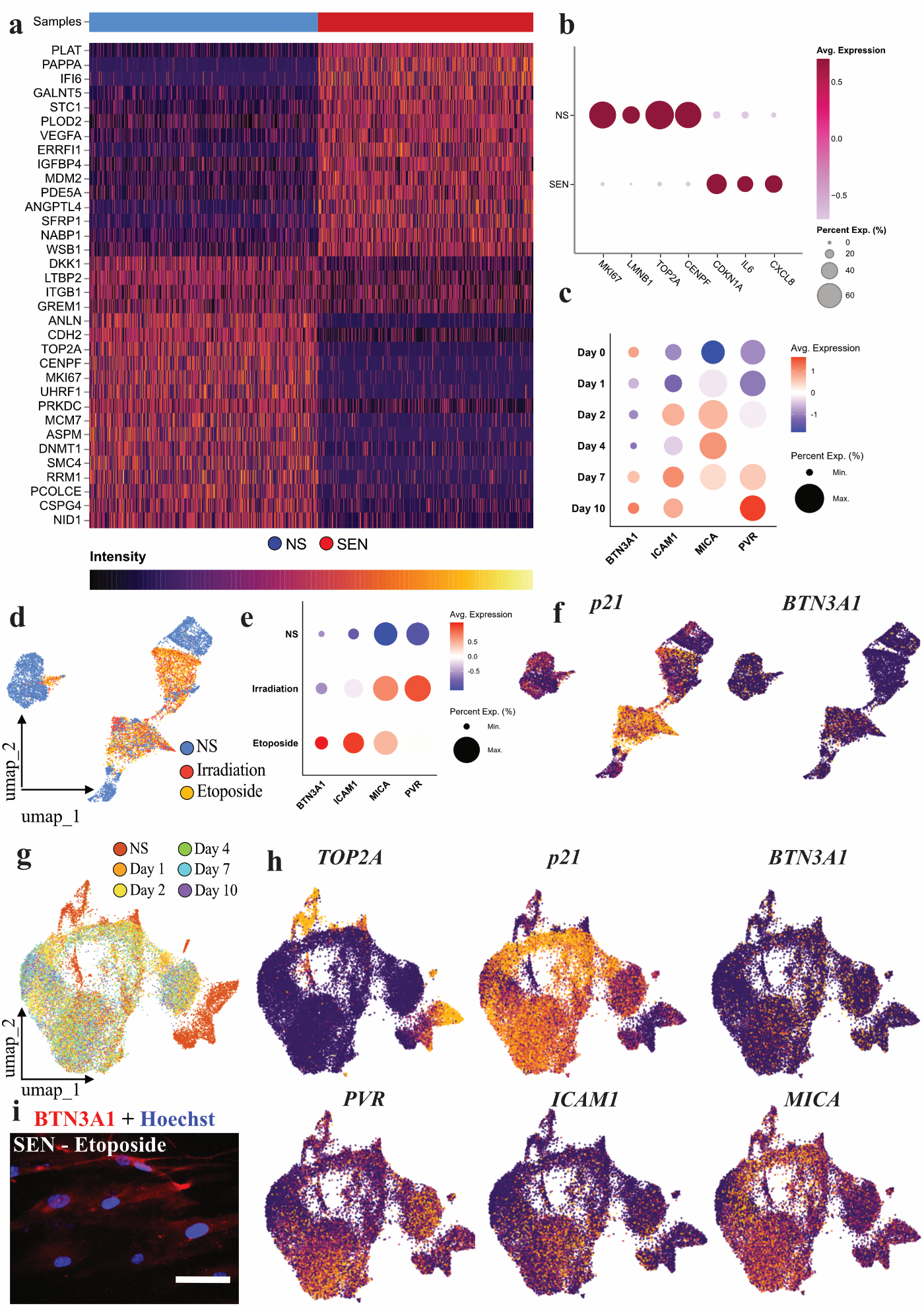

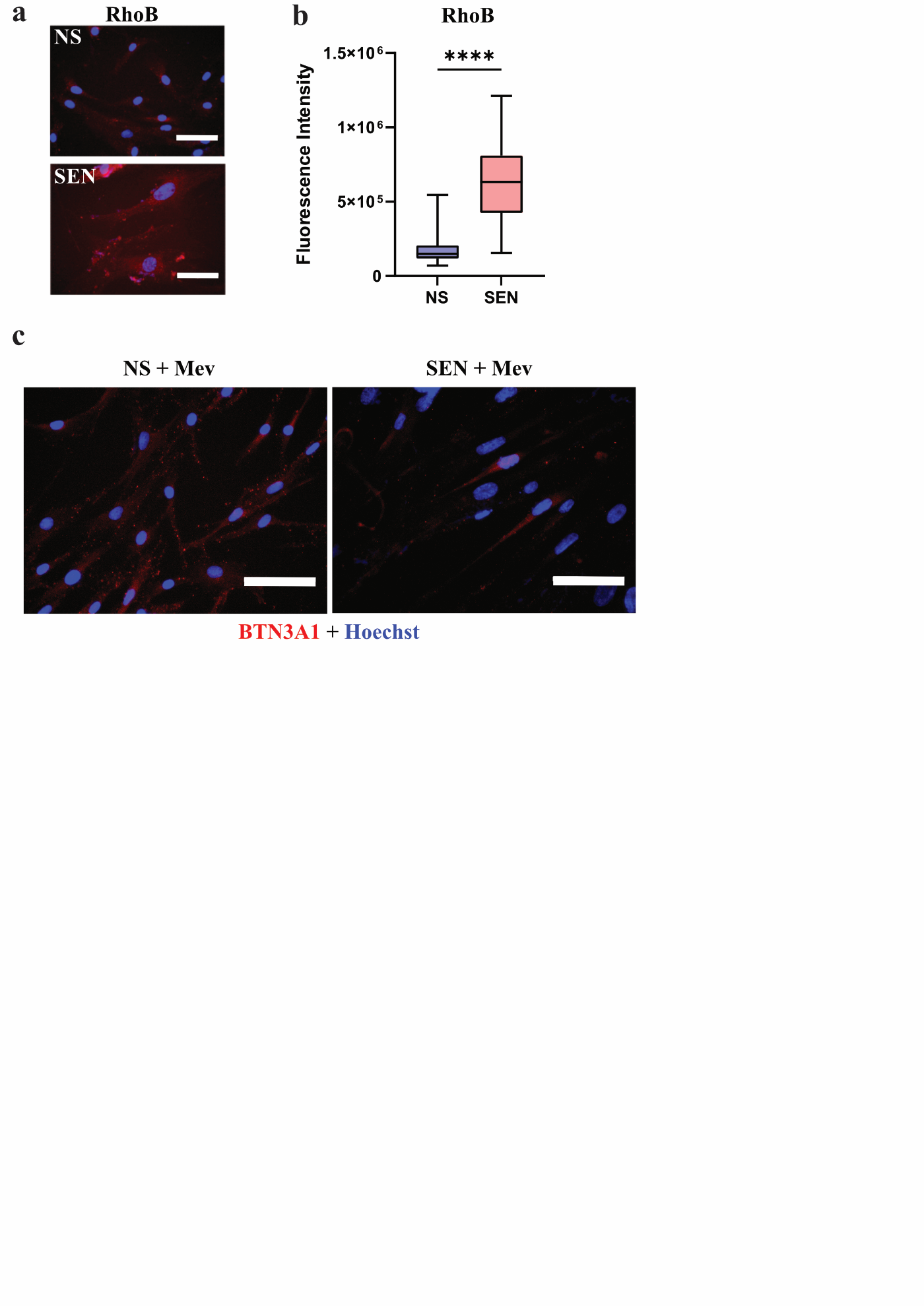

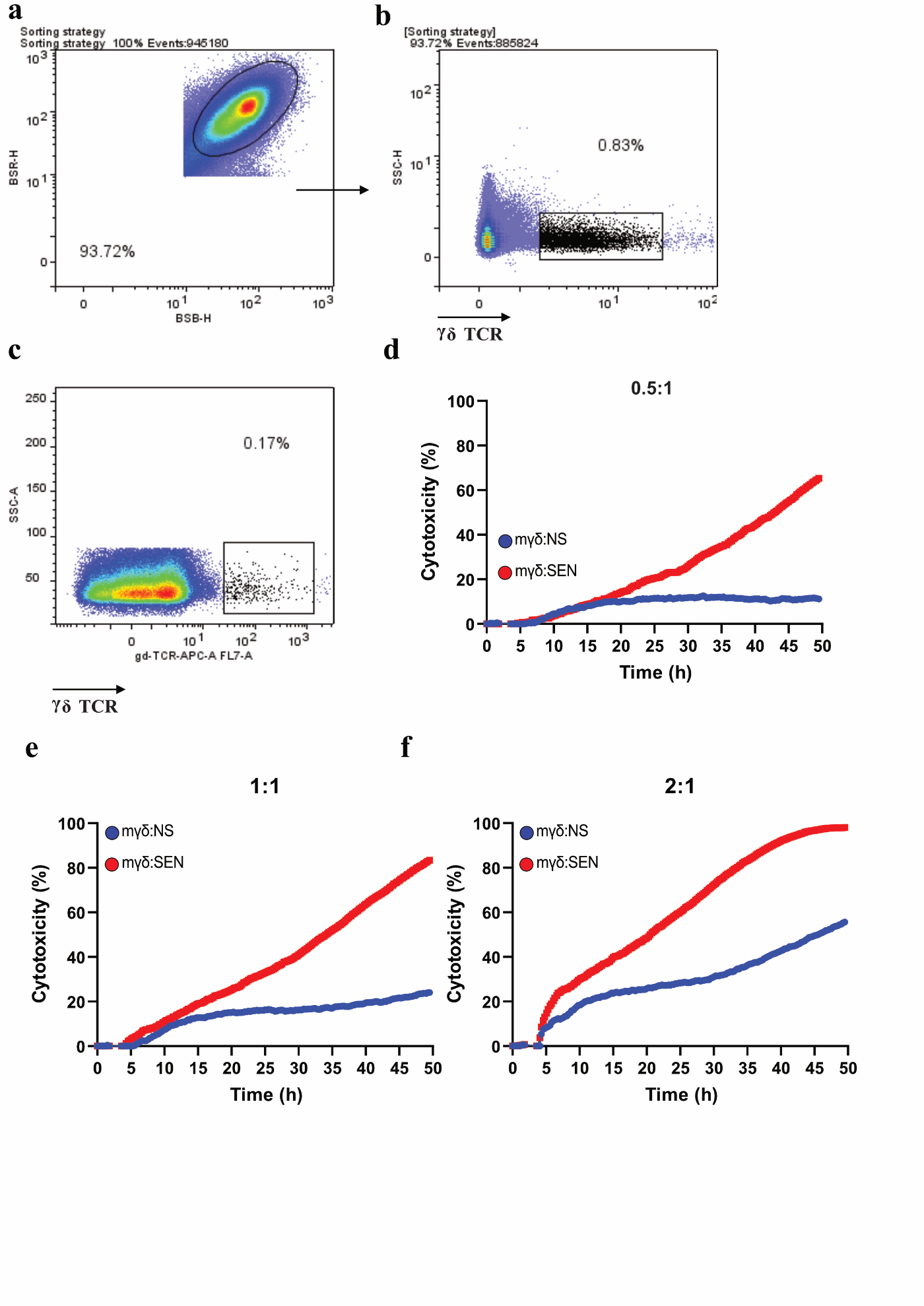

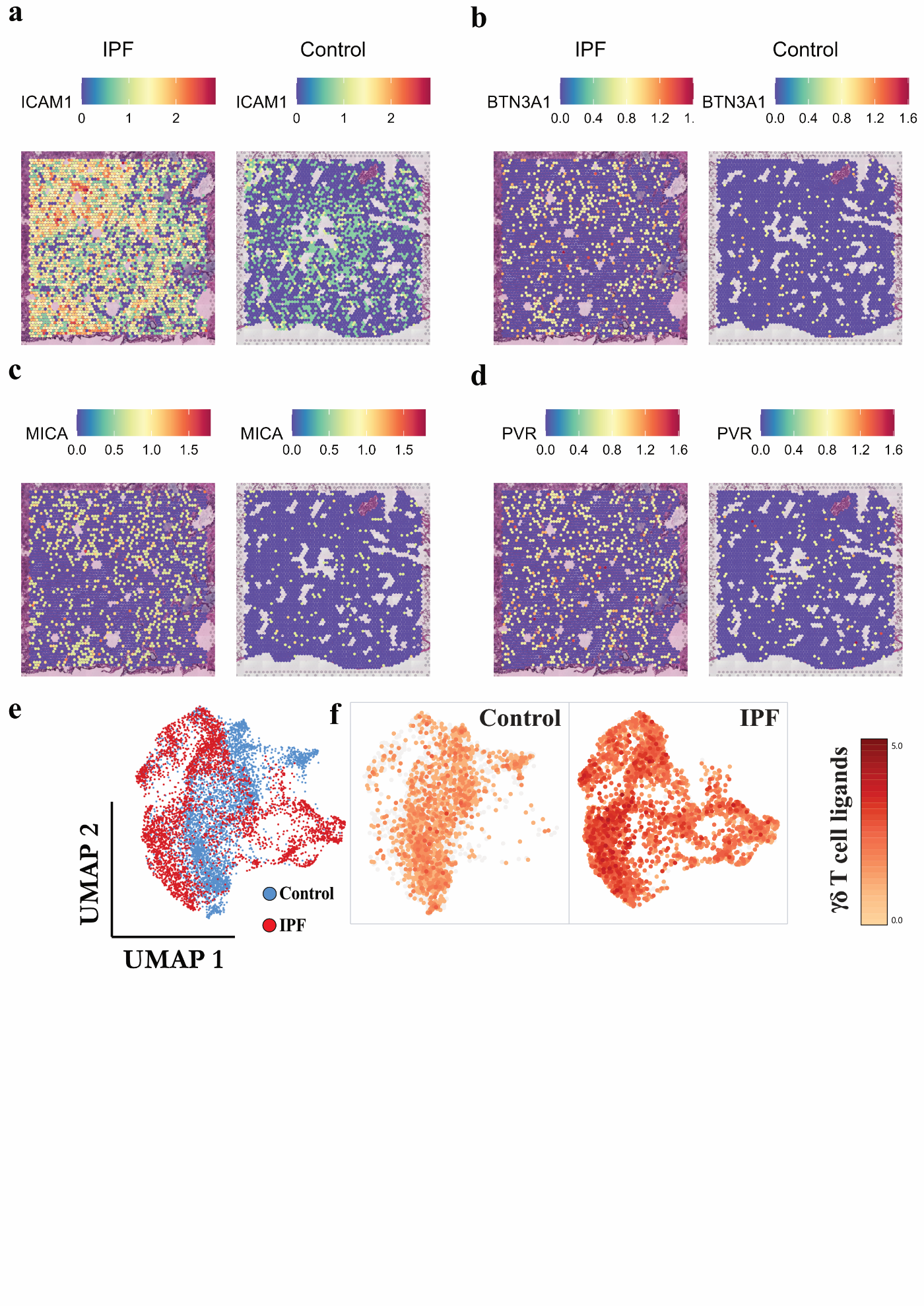

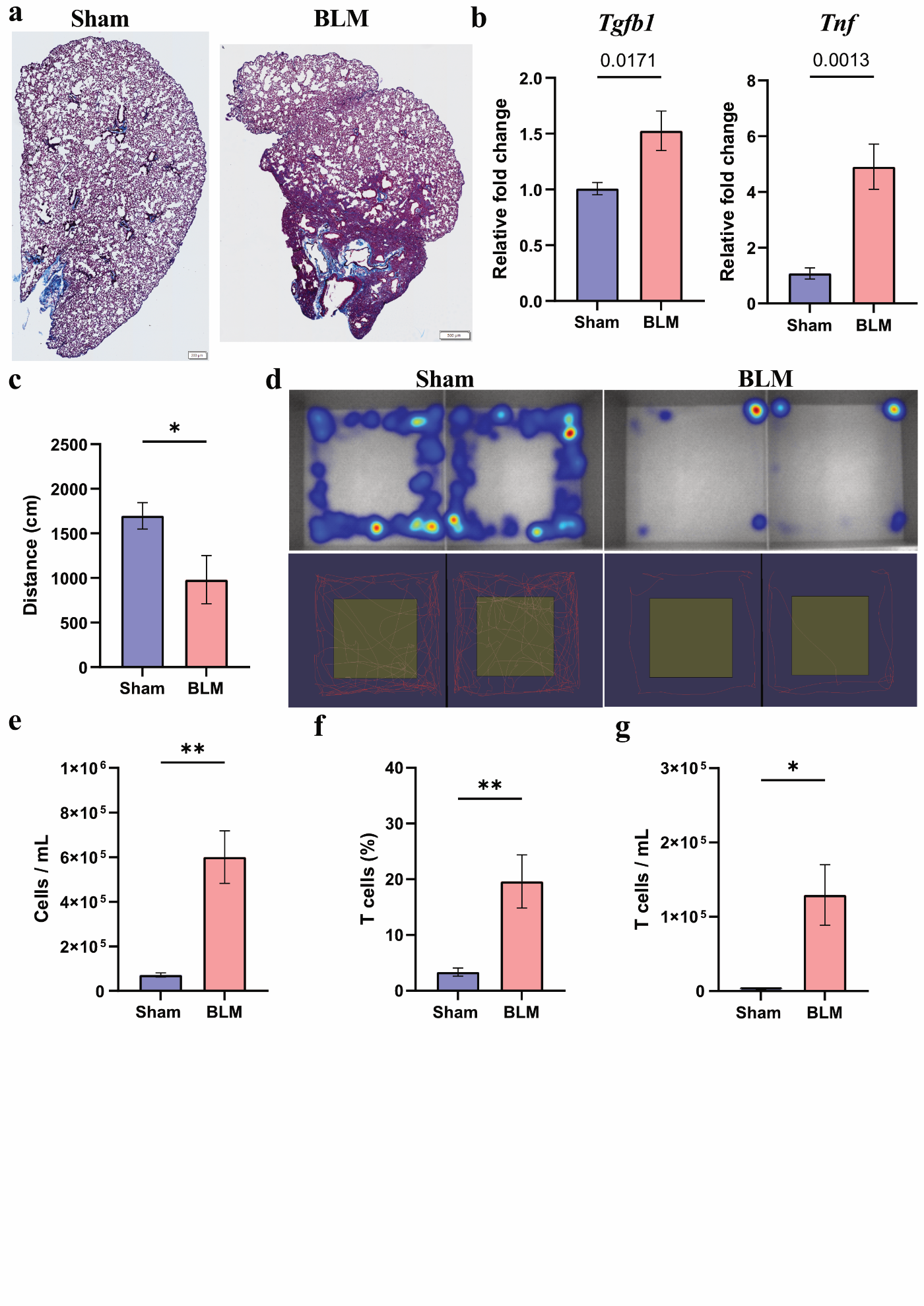

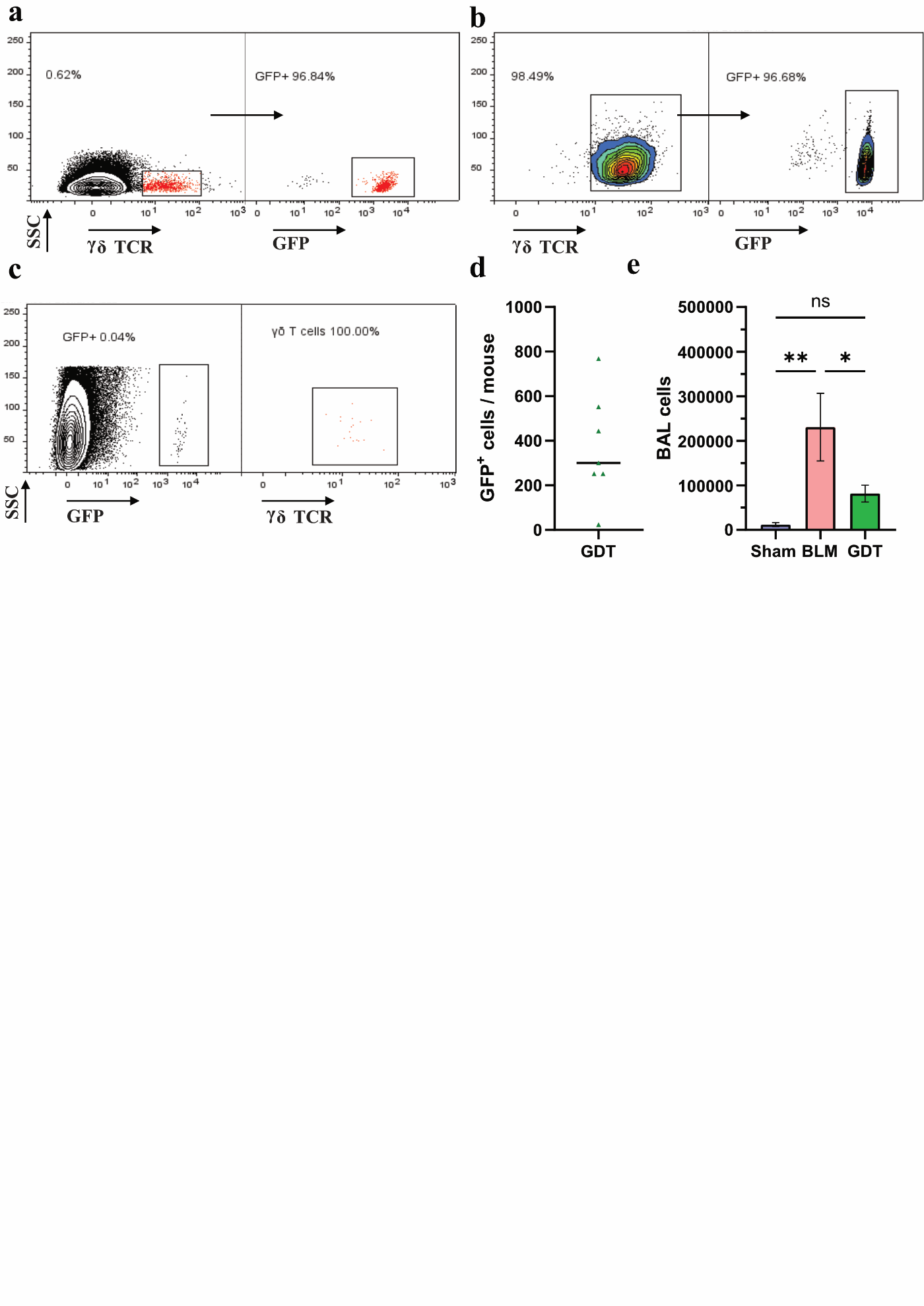

**Supplementary Figure Legends**

**Supplementary Figure 1. Characterization of naturally occurring and expanded human γδ T cells in PBMC**

(**a**) Representative gating strategy for human γδ T cells. (**b**) Flow cytometry analysis of γδ T cells representing donor-to-donor variability in the PBMC. (**c**) Stepwise flow cytometry analysis illustrating the depletion of αβ T cells and subsequent enrichment of γδ T cells. **Left panel:** Initial γδ T cells identified in the PBMC population. **Middle panel:** γδ T cells after ZOL and IL-2-mediated expansion and activation, demonstrating increased population percentage. **Right panel:** Final purified γδ T cell population, confirming successful enrichment. Representative data from experiments performed in at least three independent blood donors (n ≥ 3). (**d**, **e**) Characterization of human γδ T cell population after ZOL and IL-2 enrichment; (**d**) percentage of CD3^+^Vδ2^+^ cells after enrichment. (**e**) Percentage of CD3^+^Vδ1^-^ cells after enrichment. Representative plot from experiments performed in at least three independent donors (n ≥ 3). (**f**) γδ T cells activation and phenotype after enrichment. Percentage of CD3^+^γδTCR^+^CD56^+^ cells after ZOL and IL-2 treatment. Representative plot from experiments performed in at least three independent donors (n ≥ 3).

**Supplementary Figure 2. Specificity of human γδ T cells for senescent cell removal**

(**a**) Quantification of cells with nuclear HMGB1 after doxorubicin treatment. Data is representative of n ≥ 3 independent experiments; unpaired t-test. (**b**) Quantification of cells with ≥ 2 γH2AX foci after senescence induction. Data is representative of n ≥ 3 independent experiments; unpaired t-test. (**c**) Cytotoxicity of PBMCs to NS controls and SEN cells at E:T ratios of 0.5:1 and 1:1. Representative data from experiments performed in at least three independent donors (n ≥ 3). (**d**, **e**) Depletion of αβ T cells reduces off-target cytotoxicity (**d**) γδ T cell killing kinetics to NS controls (purple) and SEN (green) at E:T ratio of 1:1. Light red and light blue, the γδ T cell population was further depleted of αβ T cells. Representative data from experiments performed in at least three independent donors (n ≥ 3). (**e**) Endpoint cytotoxicity of γδ T cells depleted and not depleted of αβ T cells to NS controls and SEN. Representative data from experiments performed in at least three independent donors (n ≥ 3). Two-way ANOVA.

**Supplementary Figure 3. SnRNA-Seq. and scRNA-Seq. reveal upregulation of γδ T cell ligands in senescent cells irrespective of cell type or insult**

(**a**) Unsupervised heatmap of highly expressed genes in the NS (blue) and SEN (red) cells. Intensity, Z-Score. (**b**) Supervised gene expression profile of senescence biomarkers in the NS (top) and SEN (bottom) samples. (**c**) Analysis of NS and the kinetics of etoposide-induced SEN WI-38 gene expression profile for *BTN3A1*, *ICAM1*, *MICA*, and *PVR*. Data was re-analyzed from Wechter et al., 2023. (**d**) UMAP clustering of NS, etoposide-induced SEN, and irradiation-induced SEN WI-38 cells. Data was re-analyzed from Wechter et al., 2023. Number of cells after quality control = 9,670 cells; NS = 5,207 control cells; etoposide-induced SEN = 2,232 SEN; irradiation-induced SEN = 2,231. (**e**) Gene expression dot plot profile of *BTN3A1*, *ICAM1*, *MICA*, and *PVR* in NS and multiple types of SEN. Data was re-analyzed from Wechter et al., 2023. (**f**) Gene expression of *p21* and *BTN3A1* in NS and SEN WI-38 cells. Dark purple represents low/absent gene expression. Yellow represents high gene expression. Data was re-analyzed from Wechter et al., 2023. (**g**) Gene expression data re-analysis of SEN and NS WI-38 cells from Wechter et al., 2023. UMAP of NS controls and etoposide-treated WI-38 SEN at different time points. Number of cells after quality control = 22,575 cells; Day 0 NS WI-38 = 5,161 control cells. (**h**) Associated gene expression profile of TOP2A, p21, BTN3A1, PVR, ICAM1, MICA in WI-38 SEN cells. Dark orange represents higher expression; dark purple represents the absence of gene expression. (**i**) Immunofluorescent BTN3A1 staining of unpermeabilized etoposide-induced SEN. BTN3A1 (red); Hoechst (blue). Scale bar = 75 μm. Data is representative of one of ≥ 3 independent experiments.

**Supplementary Figure 4. RhoB is upregulated in senescent cells and Mevastatin treatment lowers BTN3A1 in doxorubicin-induced SEN**

(**a**) Representative images of RhoB staining in NS and SEN. Scale bar = 75 μm. Data is representative of one of 4 independent experiments. (**b**) Fluorescence intensity quantification of RhoB staining in NS and SEN. Data is representative of one of 4 independent experiments; unpaired t-test. (**c**) Representative images of CD277/BTN3A1 staining in NS and doxorubicin-induced SEN after 24 hours of Mevastatin treatment. Scale bar = 75 μm. Data is representative of one of ≥ 3 independent experiments.

**Supplementary Figure 5. Purification of mouse γδ T cells for the elimination of senescent cells**

(**a**, **b**) Representative FACS strategy to isolate mouse γδ T cells. (**a**) Splenocytes were gated based on size, and (**b**) γδ T cells were sorted based on γδ TCR expression. (**c**) Representative flow cytometry plot of γδ T cell depletion after FACS. (**d**, **e**, **f**) Mouse γδ T cell 48-hour killing kinetics to NS controls and SEN at E:T ratios of (**d**) 0.5:1, (**e**) 1:1, and (**f**) 2:1. Representative data from one experiment performed in at least three independent mouse donors (n ≥ 3).

**Supplementary Figure 6. Characterization of γδ T cell ligands in the human fibrotic lung**

(**a**, **b**, **c**, **d**) Spots expressing γδ T cell ligands are enriched in the IPF lungs. (**a**) ICAM1, (**b**) BTN3A1/CD277, (**c**) MICA, and (**d**) PVR expression in IPF and control human lung. IPF = Freshly frozen human IPF lung tissue collected during transplantation with severe tissue remodeling. Control = Healthy control lung tissue (no known lung disease) collected postmortem. No tissue remodeling was observed in the healthy controls. Data was re-analyzed from Franzén et al., 2024 (PMID: 38951642). (**e**) UMAP of IPF and control human lung. (**f**) Expression of γδ T cell ligands (ICAM1, BTN3A1/CD277, MICA, and PVR) in the IPF lungs. Orange represents low/absent γδ T cell ligands, whereas dark red represents high γδ T cell ligand abundance. Data was re-analyzed from Franzén et al., 2024 (PMID: 38951642).

**Supplementary Figure 7. Characterization of the BLM-induced fibrosis model**

(**a**) Collagen deposition in the lungs of mice treated with BLM, trichrome Masson staining. (**b**) Gene expression of *Tgfb1* and *Tnf*. RNA was isolated from the right lobes. *Gapdh* was used as housekeeping control. (**c**) Total distance (in cm) moved by sham-treated and BLM-treated mice 21 days after treatment. Spontaneous movement was measured for 10 minutes using the EthoVision platform. (**d**) Representative images of spontaneous movement of sham-treated and BLM-treated mice 21 days after treatment. On the top, movement heatmap; red areas represent more time spent in an area compared to blue. Top represents only periphery movement. On the bottom, total distance traveled by a given mouse. (**e**) Total cells per mL collected in the BAL in the sham-treated and BLM-treated mice 21 days after treatment. Singlets were used as cell readouts. (**f**, **g**) T cells are recruited after fibrotic insult. The (**f**) percentage and (**g**) number of the total mouse T cells was quantified by quantifying CD3^+^ cells from the size, singlets, and CD3^+^ gate in the BAL cells.

**Supplementary Figure 8. Characterization of the phenotype of GFP^+^** **γδ T before and after adoptive cell transfer**

(**a**, **b**) Representative flow cytometry plot of GFP^+^ γδ TCR^+^ cells. Mouse γδ T cells were isolated and enriched from the spleen of C57BL/6-Tg(UBC-GFP)30Scha/J. (**a**) The initial population of GFP^+^ γδ TCR^+^ from the spleen. (**b**) Representative flow cytometry plot of GFP^+^ γδ TCR^+^ cells after 7 days expansion with IL-2 and fluorescence-activated cell sorting purification. (**c**, **d**) GFP^+^ fraction after γδ T cell adoptive cell transfer. BAL cells were harvested 7 days after adoptive cell transfer. (**c**) Representative flow cytometry plot of the GFP^+^ fraction. GFP^+^ cells stained positive for γδ TCR. (**d**) Number of GFP^+^ cells in the γδ T cell-treated mice. BLM-treated and sham-treated mice had 0 GFP^+^ cells. (**e**) Decrease in cell number in the BAL cells. N = 6 sham-treated mice; N = 5 BLM-treated mice; N = 7 GFP^+^ γδ T cell transfer.
